# GATA4/5/6 family transcription factors are conserved determinants of cardiac versus pharyngeal mesoderm fate

**DOI:** 10.1101/2020.12.01.406140

**Authors:** Mengyi Song, Xuefei Yuan, Claudia Racioppi, Meaghan Leslie, Anastasiia Aleksandrova, Lionel Christiaen, Michael D. Wilson, Ian C. Scott

## Abstract

GATA4/5/6 transcription factors play essential, conserved roles in heart development. How these factors mediate the transition from multipotent mesoderm progenitors to a committed cardiac fate is unclear. To understand how GATA4/5/6 modulate cell fate decisions we labelled, isolated, and performed single-cell gene expression analysis on cells that express *gata5* at pre-cardiac time points spanning gastrulation to somitogenesis. We found that most mesendoderm-derived lineages had dynamic *gata5/6* expression. In the absence of Gata5/6, the population structure of mesendoderm-derived cells was dramatically altered. In addition to the expected absence of cardiac mesoderm, we observed a concomitant expansion of cranial-pharyngeal mesoderm. Functional genetic analyses in zebrafish and the invertebrate chordate *Ciona*, which possess a single GATA4/5/6 homolog, revealed an essential and cell-autonomous role for GATA4/5/6 in promoting cardiac and inhibiting pharyngeal mesoderm identity. Overall, the maintenance and repression of GATA4/5/6 activity plays a critical, evolutionarily conserved role in early development.

## Introduction

Heart development is a complex process that requires tightly controlled gene regulatory networks, which direct the correct specification, differentiation, patterning, and morphogenesis of the heart and its constituents. Transcription factors (TFs) are the core components of this regulatory network as they play critical roles in initiating cardiac specification ^1–3^. Among the earliest TFs that are expressed in the cardiac mesoderm, *GATA4, GATA5* and *GATA6* paralogs (herein referred to as *GATA4/5/6*) function at or near the top of the cardiac regulatory network hierarchy in animals ^1,4^. For example: the loss of *Gata4/6* in mice and *gata5/6* in zebrafish both lead to a heartless phenotype ^5–7^; targeted expression of a dominant negative form of the sole *GATA4/5/6* homolog in the chordate *Ciona*, *Ci-Gata-a*, results in a significant reduction of cardiac gene expression ^8^; and the loss of the *GATA4/5/6* homolog *pannier* in *Drosophila* leads to a heartless phenotype ^9^. Highlighting their essential and dosage-sensitive roles in heart development, individual heterozygous loss-of-function mutations in *GATA4/5/6* in humans have been associated with congenital heart disease ^10–12^.

Cross-species comparisons suggest that zebrafish *gata5* is the functional homolog of mammalian *Gata4* ^13–15^. Zebrafish *gata5* mutation results in greatly reduced expression of the canonical cardiac mesoderm marker *nkx2.5*, with improper migration of remaining cardiac progenitors ^14^, likely due to defects in endoderm development and morphogenesis ^16,17^. *gata6* mutation in zebrafish embryos leads to a reduction of the ventricular chamber and a dilation of the atrial chamber ^6^. Combined mutations or injection of morpholinos (MOs) targeting both *gata5* and *gata6* completely blocks cardiogenesis, with an absence of the *nkx2.*5-expressing cardiac progenitors ^5,6^. Despite these insights, how *GATA4/5/6* family members facilitate the early separation of the cardiac lineage remains unclear.

Zebrafish *gata5* and mouse *Gata4* are amongst the first TFs expressed in precardiac mesoderm ^14,18^. Lineage tracing, clonal analyses, and single-cell transcriptome profiling suggest that cardiac progenitors with distinct potential and transcriptional profiles are present as early as the onset of gastrulation ^19–24^. At least two distinct cardiac progenitor populations, the first heart field (FHF) and second heart field (SHF), arise during gastrulation, with the SHF being composed of late-differentiating progenitors that add to the developing heart tube derived from the FHF. Although in mouse *Gata4/6* seem to not be required for early specification of the SHF ^7^, they later function together with Hedgehog signaling to regulate outflow tract development ^25^. It therefore remains to be resolved at which stages the *GATA4/5/6* family of transcription factors is essential for cardiogenesis, and which processes they regulate.

The broad expression and the pioneering function of GATA family TFs ^26^ are consistent with their central role in initiating diverse developmental programs. *GATA4/5/6* transcripts are broadly localized in the mesendoderm during vertebrate gastrulation ^14,18,27–29^, well before expression of the canonical cardiac and pharyngeal mesoderm marker *NKX2.5* is initiated. *In vivo*, overexpression of *Gata4* (mouse) or *gata5* (zebrafish), in concert with other factors, promotes cardiac gene expression programs and phenotypes ^30,31^. However, *GATA4/5/6* do not solely regulate cardiogenesis, with mouse *Gata4* ^32^ and zebrafish *gata5* ^14^ playing essential roles in endoderm development. *Gata5/6*-deficient zebrafish embryos also display other mesodermal defects including reduction of anterior haemangioblast and fin bud progenitors ^5,33^. Given their potent pro-cardiac activity, it remains to be understood how *GATA4/5/6* family members facilitate the early divergence of cardiac and non-cardiac lineages.

To investigate how *gata5/6* regulate the early divergence of the cardiac lineage, we used a reporter line to isolate *gata5*-GFP positive cells from zebrafish embryos at stages spanning early gastrulation to early segmentation stages and subjected them to single-cell mRNA sequencing (scRNA-seq). We found dynamic, lineage-specific regulation of *gata5/6* transcripts, with *gata5/6* expression being maintained in the cardiac lineage while being gradually turned off in the pharyngeal mesoderm. Single-cell RNA-seq analysis of wild-type (WT) and Gata5/6 knockdown embryos revealed both expected (loss of cardiac mesoderm and reduced endoderm) and unexpected (reduced pronephric mesoderm, expanded erythroid progenitors and pharyngeal mesoderm) changes in cell lineage composition. Zebrafish transplantation experiments showed that Gata5/6 knockdown promotes pharyngeal mesoderm fate in a cell-autonomous manner.

Finally, targeted over-expression and CRISPR-mediated loss of the single *GATA4/5/6* homolog in cardiopharyngeal progenitors of the tunicate *Ciona* indicated that sustained *GATA4/5/6* expression is a conserved mechanism that promotes cardiac fate specification, in part by preventing ectopic activation of the pharyngeal muscle program. Overall, we suggest that *GATA4/5/6* family members are central regulators of a conserved mechanism underlying the divergence of cardiac and pharyngeal fates.

## Results

### Dynamic expression of *gata5* in a broad range of mesoderm lineages

The heartless phenotype in zebrafish *gata5/6* mutants and morphants can first be detected at the bilateral heart field stage, with loss of *nkx2.*5, the earliest apparent definitive marker of cardiac mesoderm. However, the expression of *gata5* and *gata6* can be detected prior to the onset of gastrulation, broadly within the mesendoderm and yolk syncytial layer (YSL), and become restricted to the anterior lateral plate mesoderm (ALPM) by 13 hours post-fertilization (hpf, Supplementary Fig. 1.1a-d) ^14,29^. This suggests that *gata5* and *gata6* are required for the early specification of cardiac progenitor cells, likely during or immediately after gastrulation. We therefore sought to profile how *gata5/6* facilitate the early segregation of the cardiac lineage. We utilized the *TgBAC (gata5:EGFP)*^*pd25*^ line ^34^ that faithfully recapitulated the endogenous expression of *gata5* (Supplementary Fig. 1.1). Importantly, due to the perdurance of GFP and the rapid development of early zebrafish embryos, this approach allowed us to capture all cells that expressed *gata5* prior to 13 hpf. We first conducted scRNA-seq on *gata5:*GFP+ cells at times spanning early to late gastrulation (shield: 6 hpf, 75% epiboly: 8 hpf, and tailbud: 10 hpf), and early somite stages (8 somite stage (ss): 13 hpf) (Fig. 1a). As *gata5:*GFP+ cells make up 6-15% of the embryo at the stages that we profiled, we obtained scRNA-seq data for 901-1,985 cells for each stage using the 10X Genomics platform, in theory reaching 2-3 times more coverage of the *gata5*:GFP+ populations compared to a published whole embryo single-cell dataset ^35^. After stringent quality control, 5,358 cells were used for downstream analyses (Supplementary Table 1).

**Fig. 1:**
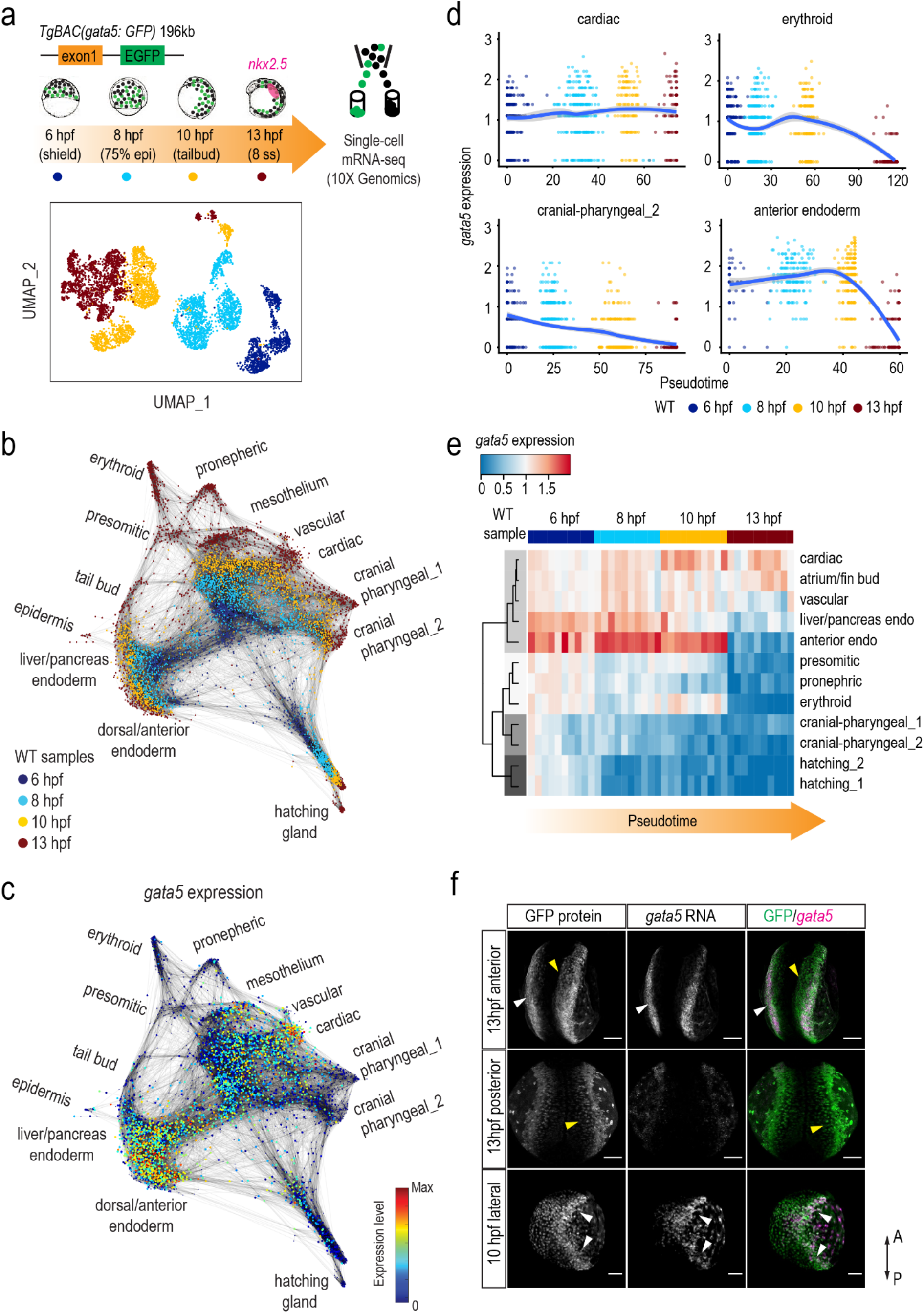
Dynamic regulation of *gata5* in various mesendoderm lineages. **a**, Schematic representation of the scRNA-seq experimental design and UMAP visualization of *gata5*:GFP+ cells collected at 4 developmental time points. *TgBAC(gata5:GFP)* contains a 196 kb genomic region spanning the *gata5* locus, with the *GFP* sequence inserted after the first exon of *gata5*. **b**, Force-directed graph showing the connection between all single cells from the four WT samples. Nodes were colored by developmental time points. Edges link nodes with their neighboring cells. For each cell, up to 20 mutual nearest neighbor edges within or between time points are retained. **c**, Expression of *gata5* projected on the force-directed graph. **d**, Pseudotemporal expression profiles of *gata5* along the developmental trajectories of representative lineages (cardiac, pharyngeal_2, erythroid, anterior endoderm). x-axis, pseudotime; y-axis, normalized log expression level. The smoothed lines show conditional means after local polynomial regression fitting (LOESS method) and shaded areas indicate standard errors. **e**, Heatmap showing *gata5* expression along the developmental trajectory of major lineages. Pseudotime was scaled across all lineages for each stage. Each column represents a quantile bin of the scaled pseudotime and each row a lineage. *gata5* expression mean was calculated for each pseudotime bin for each lineage. The row dendrogram shows the unsupervised hierarchical clustering of each lineage based on *gata5* expression dynamics. **f**, FISH against *gata5* transcripts and immunostaining against GFP at 10 hpf and 13 hpf. White arrows show cells positive for both *gata5* transcripts and GFP protein; yellow arrows indicate cells positive for GFP protein only. A: anterior, P: posterior. All scale bars represent 100 μm.

We first performed graph-based clustering to examine the cellular heterogeneity of *gata5:*GFP+ cells at each stage ^36,37^. Consistent with previous reports, we observed an increasing complexity of mesodermal and endodermal lineages from 6 to 13 hpf ^35,38^, with a divergence of cells evident as early as 6 hpf (Supplementary Figs. 1.2-1.5). At 10 hpf expression of *ttn.2*, which encodes the cardiomyocyte-specific sarcomeric component Titin, was evident in an apparent cardiac mesoderm group that co-expressed other genes associated with cardiogenesis (*nkx2.7, gata6, foxh1*) (cluster 7, Supplementary Fig. 1.4c), suggesting that expression signatures of cardiac commitment are evident by the end of gastrulation. Cellular heterogeneity was greatly increased at 13 hpf, with the emergence of 16 discernible clusters (Supplementary Fig. 1.5).

Consistent with known roles of *gata5/6* ^5,14^, we detected cardiac progenitors (cluster 9: *nkx2.5*, *mef2ca*, *mef2cb*) and two distinct endoderm populations (cluster 5: *foxa2*, *nkx2.7*; cluster 12: *onecut1*, *angptl3*). Using established marker genes we further identified unexpected *gata5-* labeled cell types at 13 hpf: erythroid (cluster 1: *gata1a, cldng, gfi1aa*), endothelial (cluster 11: *etv2, fli1b, kdrl*), pronephric (cluster 2: *acy3.2, hnf1bb, foxj1a*), presomitic (cluster 10: *tbx16*, *bambia*), somitic (cluster 16: *myod1, ttn.1, myf5*), and two cranial-pharyngeal (cluster 3: *twist1b*, *thbs3b*; cluster 7: *pitx3, foxf2a*) populations. We also observed a likely mesothelium progenitor population (cluster7: *hand2*, *cxcl12a, cdx4*) ^39^ (Supplementary Fig. 1.5).

To further interpret the developmental progression of GFP+ cells from 6 to 13 hpf, we employed a graph-based method to ‘stitch’ cells with similar gene expression from consecutive time points and visualized using a force-directed layout ^35^. We observed three major branches representing mesoderm, endoderm, and hatching gland starting from 6 hpf. At 13 hpf (red dots), branching tips representing distinct cell types are evident, including all major lineages that we observed in the clustering analysis (Fig. 1b and Supplementary Fig. 1.6). Taken together, this indicated a broad spectrum of lineages that arise from early *gata5*-expressing progenitors.

As the role of *gata5* has not been linked to the specification of many of the lineages identified at 13 hpf ^32,40^, we next examined *gata5* expression in our scRNA-seq data (Fig. 1c). In most lineages at 13 hpf (branching tips), including hatching gland, pharyngeal mesoderm, and posterior mesoderm derivatives (erythroid, presomitic, and pronephric progenitors), *gata5* was no longer expressed, with high levels of *gata5* only evident in cardiac, vascular, and mesothelium progenitors (Fig. 1c). To better delineate the temporal expression dynamics of *gata5*, we established pseudotime developmental trajectories for each distinct lineage (see Methods for details). The pseudotime trajectory corresponded well with actual developmental time points, with *gata5* expression being regulated in distinctive manners in different trajectories (Fig. 1d and Supplementary Fig. 1.7). We next aligned the pseudotime of all trajectories to better compare the regulation of *gata5* in different differentiation paths. Though the expression of *gata5* was relatively robust at the beginning of all lineage trajectories, its expression was only maintained in the cardiac, atrium/fin bud, and vascular progenitors while being down-regulated over 8 to 13 hpf in most other lineages. Although expression of *gata6* was relatively sparse, as compared to *gata5*, we observed that *gata5* and *gata6* shared similar expression dynamics in almost all trajectories (Fig. 1e and Supplementary Fig. 1.7).

The observation of 13 hpf populations that lack detectable *gata5* expression was consistent with the perdurance of GFP protein, which exceeds the 7 hours of developmental time we examined. To confirm this, we conducted fluorescent RNA *in-situ* hybridization (FISH) against *gata5* transcripts and immunofluorescent staining against GFP protein at 10 and 13 hpf. Consistent with our trajectory analysis, cells actively expressing *gata5* were largely located at the ALPM and YSL at 13 hpf, whereas GFP+ cells that were located more medially or posteriorly did not express detectable *gata5* transcripts (Fig. 1f). Taken together, our transgenic system captured cells that expressed *gata5* prior to 13 hpf, with regulation of *gata5* and *gata6* being highly dynamic within these lineages. Both the maintenance and inhibition of *gata5/6* expression may therefore be important events in the differentiation of these lineages.

### Loss of Gata5/6 lead to changes in mesodermal cell compositions

To directly investigate roles for *gata5/6* in early lineage commitment, we conducted scRNA-seq on 13 hpf *gata5*:EGFP*+* cells collected from Gata5/6 morphants. We first combined the 13 hpf WT and morphant scRNA-seq data and followed a similar clustering analysis pipeline as above (Fig. 2a, b, and Supplementary Fig. 2a, b). We observed a slight increase in cluster numbers, likely due to the boost of analytic power obtained from doubling the number of cells (Fig. 2b and Supplementary Fig. 1.4). Overall, we detected very similar cell types in both datasets, with marker genes of most clusters shared between WT and Gata5/6 morphant cells (Fig. 2a-c and Supplementary Fig. 1.4). A notable exception was cardiac progenitors (cluster 10), which were largely absent in morphants (Fig. 2c, d). While cell identities were highly comparable between WT and Gata5/6 morphants, we observed significant changes in the overall proportions of many lineages. Aside from the absence of the cardiac lineage, there was a significant expansion of cranial-pharyngeal mesoderm (clusters 1, 2, 5, 13) and one erythroid progenitor population (cluster 4), with reduction of several posterior mesoderm clusters including those with pronephric and presomitic mesoderm signatures (Fig. 2d). These lineage proportion changes remained evident when the cardiac cluster was removed from the WT data (Supplementary Fig. 2c). Expansion and reduction of cell types not currently thought to be dependent on Gata5/6 function suggested that regulation of *gata5/6* expression may play an unappreciated role in multiple, perhaps antagonistic, fate decisions.

**Fig. 2:**
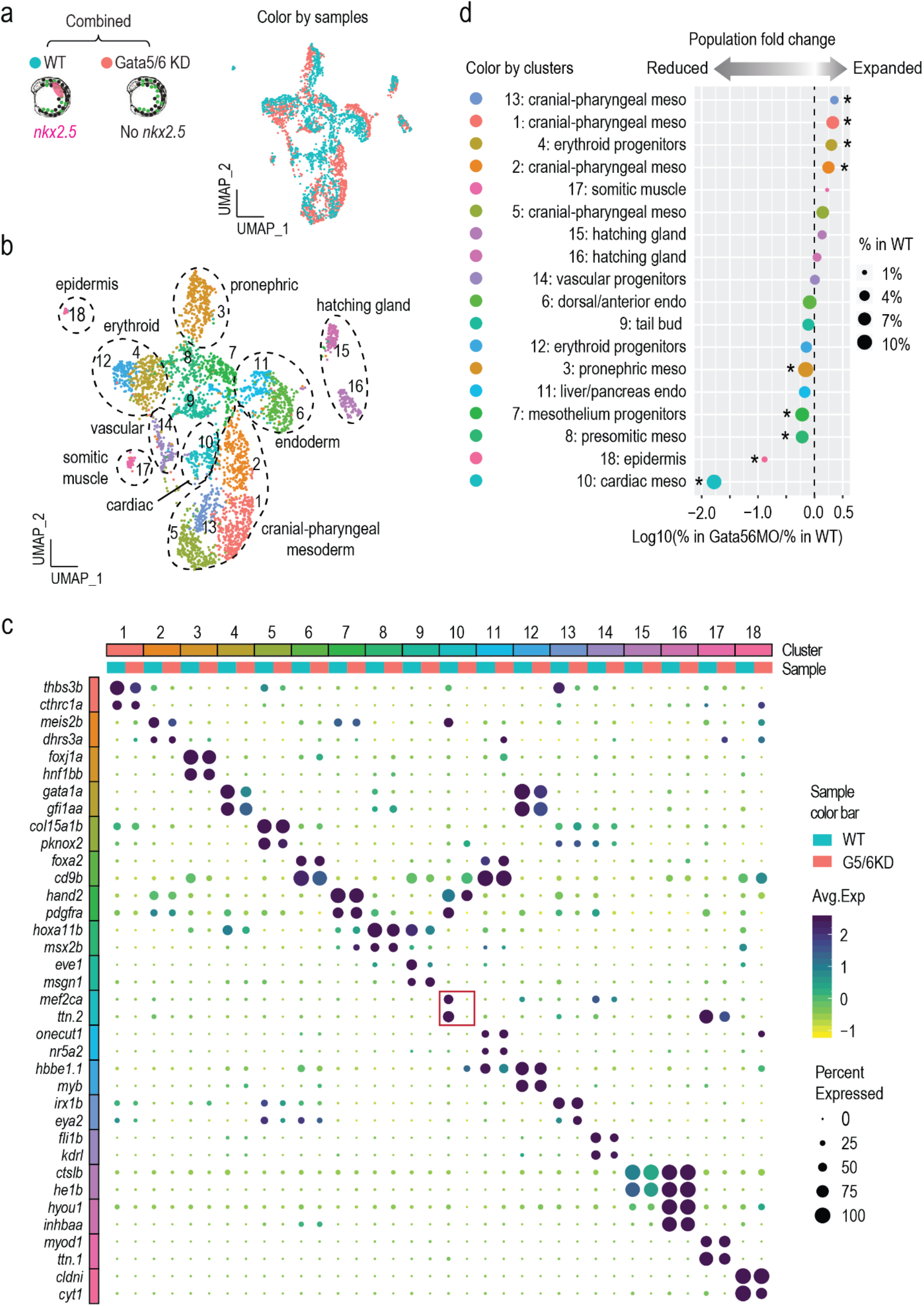
Loss of Gata5/6 leads to expansion and reduction of multiple mesoderm lineages. **a, b**, UMAP visualization of merged single-cell datasets from 13 hpf WT and Gata5/6 KD samples, colored by sample (**a**) or cluster IDs (**b**). **c**, Dot plot showing the mean expression levels (color) of markers genes and the percentages of cells in which marker genes are expressed (size) in each cluster, with WT and Gata5/6 KD cells plotted separately. Red square indicates the marker gene expression in the cardiac lineage. **d**. Cell composition changes of each cluster between Gata5/6 KD and WT samples. Asterisks indicate significant differences (Fisher’s exact test, Bonferroni correction, adjusted *p*-value < 0.05). Dot sizes show the percentage of each cluster within the whole WT population.

### Molecularly distinct cardiac and pharyngeal subtypes are evident at early somitogenesis

The most striking effect observed in the Gata5/6 morphant scRNA-seq data was the reciprocal decrease in cardiac and increase in pharyngeal mesoderm populations (Fig. 2d). This observation relates to a recently demonstrated common cardiopharyngeal field origin of cardiac and pharyngeal mesoderm ^41–43^. To further investigate the heterogeneity of the cardiac and pharyngeal clusters, we sub-clustered these cell populations (cluster1, 2, 5, 10, 13). We identified nine molecularly distinct cardiac and pharyngeal clusters, including three cardiac (sub-clusters S4, S8, S9) and six cranial-pharyngeal (sub-cluster S1, S2, S3, S5, S6, S7) mesoderm populations (Fig. 3a, Supplementary Fig. 3a). In addition to a clear cardiac population (sub-cluster S4: *mef2cb*, *hand2*), we also identified clusters expressing markers of putative atrium/pectoral fin bud (sub-cluster S8: *nr2f1a, hoxb5b*) ^44,45^ as well as SHF progenitors (sub-cluster S9: *fgf10a, fgf8a*) ^46–48^ (Fig.3b).

**Fig. 3:**
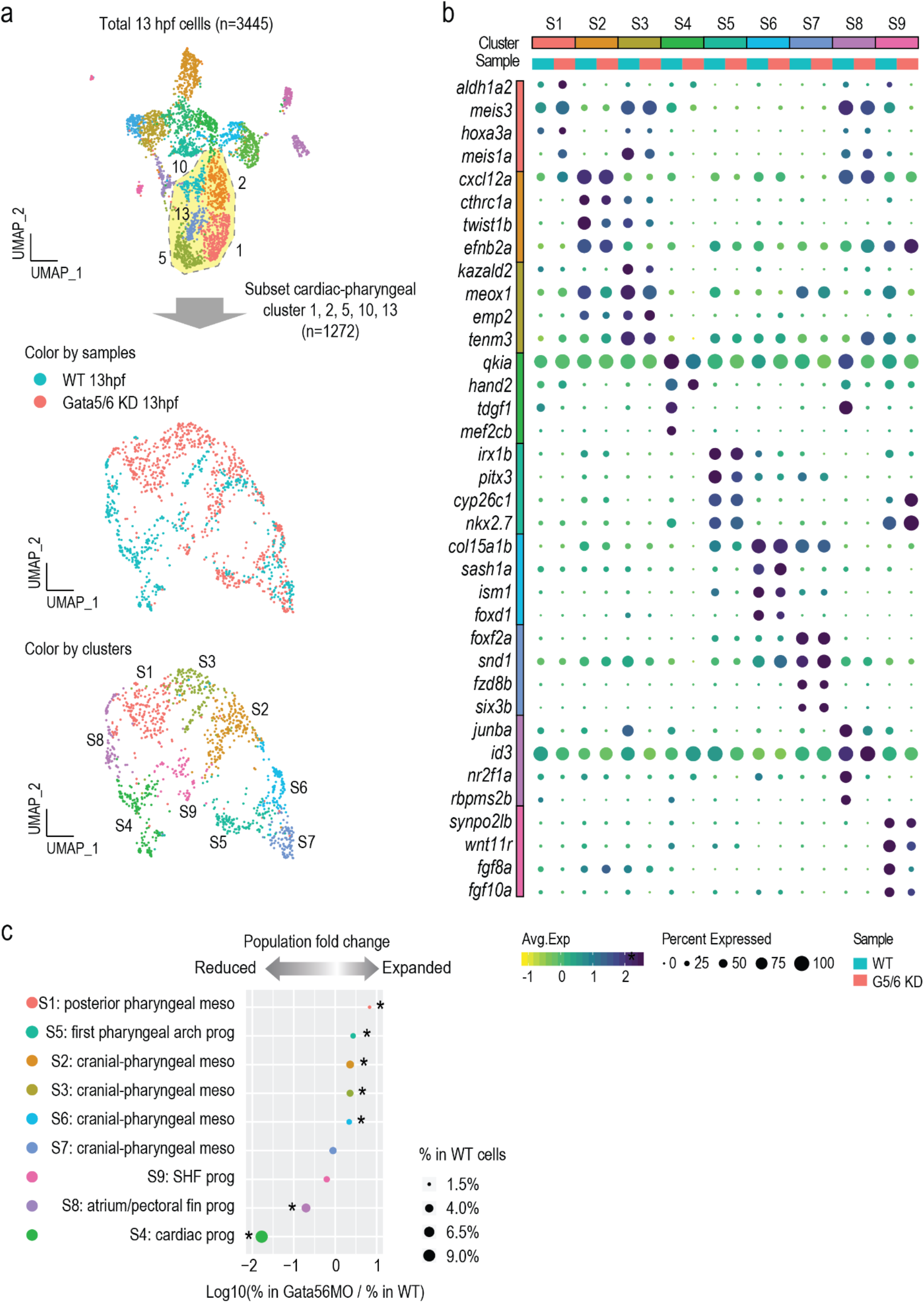
High-resolution clustering analysis of the cardiac and pharyngeal mesoderm populations. **a**, Sub-clustering analysis of the cardiac and pharyngeal mesoderm lineages. Cluster 1, 2, 5, 13, 14 were subset from the combined 13 hpf dataset shown in Fig. 2 and reanalyzed. New clustering results are visualized by UMAPs that are colored by samples (top) and by cluster IDs (bottom). **b**, Dot plot showing the mean expression levels (color) of marker genes and the percentages of cells in which marker genes are expressed (size) in each cluster, with WT and Gata5/6 KD cells plotted separately. **c**, Cell composition changes of each cluster (after sub-clustering) between Gata5/6 KD and WT samples. Asterisks indicate significant differences (Fisher’s exact test, Bonferroni correction, adjusted *p*-value < 0.05). Dot sizes show the percentage of each cluster within the whole WT population (total 13 hpf WT cells).

While SHF progenitors have been identified in several mouse single-cell studies at early stages (E7.25) ^21,22^, potential molecular signatures of the SHF at 13 hpf are largely unknown in zebrafish. Supporting that we identified zebrafish SHF cell populations, the 62 zebrafish marker genes of the SHF-like sub-cluster (S9) were enriched for regulation of SHF cardioblast proliferation (GO:0003266, FDR = 1.45e-02 ^49,50^. We next asked if the mouse orthologs of the 62 zebrafish marker genes found in our putative SHF-like sub-cluster (S9) were also found in SHF progenitors cell populations identified by comprehensive mouse single-cell studies at early stages (E7.75 and E8.25) ^51^. Indeed, among all our cardiopharyngeal sub-clusters (Supplementary Fig. 3b), the SHF-like sub-cluster (S9) overlapped the most (12/62, 19.4%) with the genes defining the mouse anterior SHF (n=212, corresponding to 221 orthologs in zebrafish).

To spatially map the six cranial-pharyngeal sub-clusters relative to the cardiac sub-clusters, we first conducted chromogenic RNA *in-situ* hybridization (ISH) for marker genes of each cardiac and pharyngeal sub-cluster (Fig. 4 a, b, d, e and Supplementary Fig. 4a-d). The putative *fgf8a+* SHF progenitor population occupied a more anterior and medial domain compared to the other cardiac progenitors at 13hpf (Supplementary Fig. 4a), as expected from previous fate-mapping studies ^52^. Moreover, by performing double FISH coupled with GFP immunostaining in *gata5:EGFP* embryos, we confirmed that the putative pharyngeal arch one progenitors had low GFP expression and were located medially to high GFP cells that were likely to be cardiovascular progenitors (Supplementary Fig. 4c). This further supports that *gata5/gfp* transcription was selectively turned off in the cranial-pharyngeal populations. Overall, we were able to spatially and molecularly resolve distinct cardiac and cranial-pharyngeal populations at early somite stages (Supplementary Fig. 4d), in agreement with previous fate-mapping work showing very early divergence of pharyngeal mesoderm sub-populations ^53^ and atrial/ventricular progenitors ^54^.

**Fig. 4.**
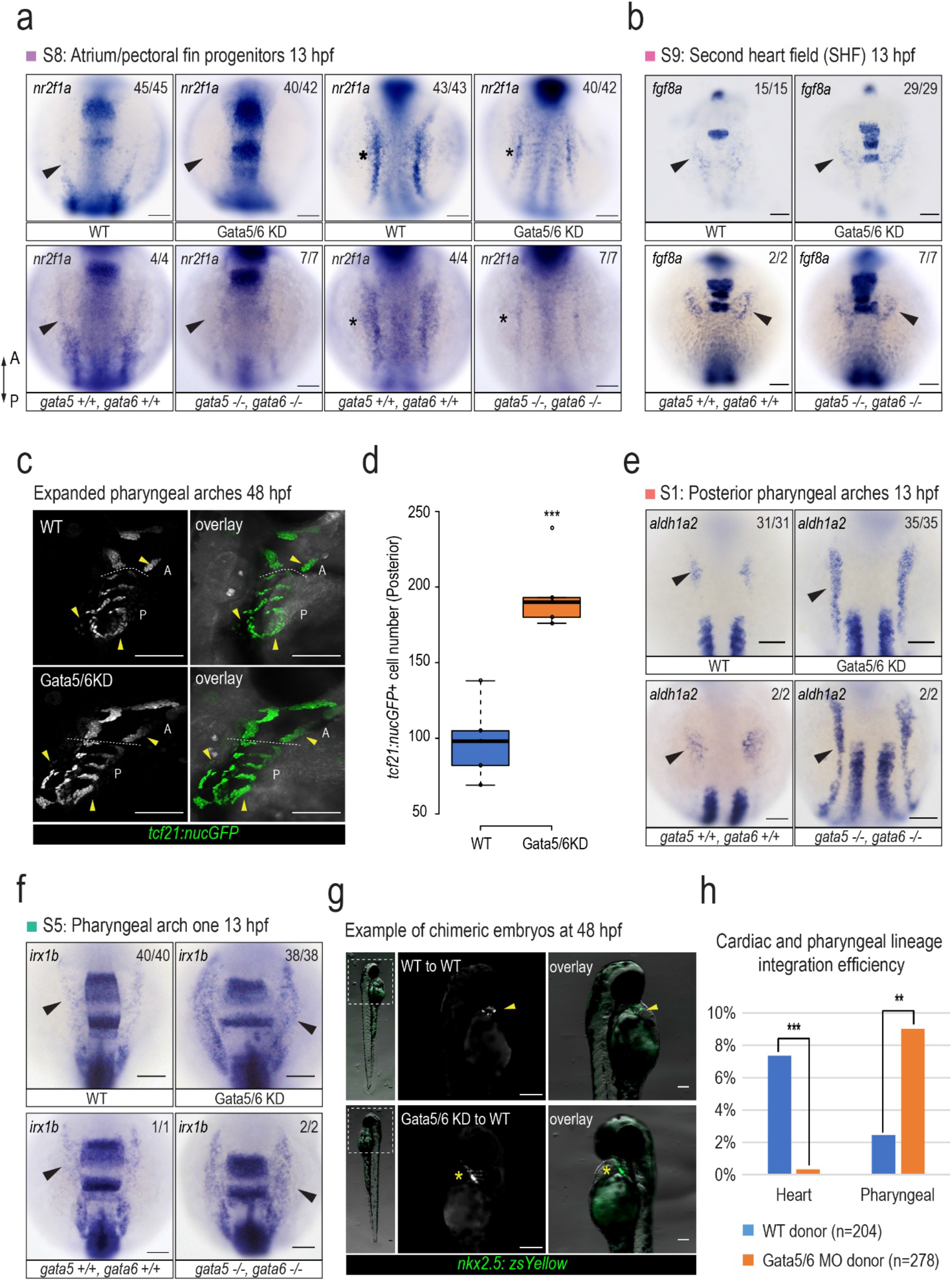
Specification defects of the cardiac and pharyngeal mesoderm upon Gata5/6 loss. **a**, RNA ISH against an atrial/pectoral fin progenitor marker gene *nr2f1a* (S8). Arrowheads indicate absent expression of *nr2f1a* in the ALPM region. Stars denote reduced expression in the putative pectoral fin progenitors. A: anterior, P: posterior **b**, RNA ISH against SHF marker gene *fgf8a* (S9). Arrowheads show that *fgf8a* expression in anterior lateral plate mesoderm is unaffected upon loss of *gata5/6.* **c**, Confocal images of *tcf21+* pharyngeal muscle cells in control and Gata5/6 morphants at 48 hpf. Arrowheads indicate expanded anterior and posterior pharyngeal arches upon Gata5/6 loss. A: anterior pharyngeal arches. P: posterior pharyngeal arches. Yellow arrowheads indicate the expanded regions. **d**, Quantification of the number of cells with *tcf21: nucGFP* activity within the posterior pharyngeal arches (arches 3-7) at 48 hpf. T-test was used to determine statistical significance. **e-f**, RNA ISH against pharyngeal markers *aldh1a2* (S1: posterior pharyngeal arches) and *irx1b* (S5: pharyngeal arch one). Arrowheads indicate their expanded expression domains in *gata5/6* deficient embryos compared to controls at 13 hpf. **g**, Two representative chimeric embryos when *Tg (nkx2.5:zsYellow)* transgenics were used as donors in transplant experiments, with arrowheads highlighting cells committed to a cardiac fate and stars indicating a pharyngeal fate. **h**, Quantification of integration efficiency. Fisher’s exact test was used to determine statistical significance. ***: p-value < 0.001, **: p-value < 0.01. All scale bars represent 100 μm.

### Gata5/6 regulates the balance and composition of cardiac versus pharyngeal fates

To examine the roles of *gata5/6* in the cardiac and pharyngeal sub-clusters we performed a cell composition analysis. This revealed distinct effects of Gata5/6 loss on different cardiac sub-populations, with the pan-cardiac (sub-cluster S4) and atrium/fin (sub-cluster S8) progenitors greatly reduced, while the SHF progenitors (sub-cluster S9) were not significantly affected (Fig. 3c). To validate these observations, we first used RNA ISH to examine the expression of *tbx5a*, a marker for cardiac and fin bud progenitors ^55^. The expression of *tbx5a* was no longer detected in cardiomyocytes while being reduced in the developing pectoral fin bud at 24 hpf upon Gata5/6 loss (Supplementary Fig. 4e). We next examined *nr2f1a*, which was expressed in both the ALPM and a more posterior domain at 13 hpf, likely representing atrial and pectoral fin bud progenitors, respectively ^44^. The ALPM expression of *nr2f1a* was completely depleted while its expression in the more posterior domain was greatly reduced in Gata5/6 deficient embryos, which likely represented impaired specification of the atrial/pectoral fin bud progenitors (Fig. 4a). Next, to confirm the existence of a SHF-like population upon loss of Gata5/6, we analyzed the expression of *tbx1* and *fgf8a* and found their ALPM expression remained largely unaltered in Gata5/6 morphants (Fig. 4b and Supplementary Fig. 4f). This suggests that SHF progenitors may still be specified upon loss of Gata5/6, indicating that specification of different cardiac progenitor subtypes have distinct requirements for *gata5/6*.

Cell composition analysis showed significant expansion in five of the six cranial-pharyngeal mesoderm sub-clusters upon loss of Gata5/6 (Fig. 3c). To characterize this further, we first injected Gata5/6 MO into the *TgBAC(tcf21: NLS-EGFP)*^*pd41*^ transgenic background, which marks the developing pharyngeal muscle progenitors ^56^. We observed an expansion of *tcf21+* pharyngeal arches (Fig. 4c), in particular the posterior arches, in Gata5/6 deficient embryos at 48 hpf (*p*-value < 0.0004; Fig. 4d). At 13 hpf, the *aldh1a2*-expressing posterior pharyngeal arches (sub-cluster S1) ^57–59^ showed the strongest expansion in Gata5/6 morphants at 13 hpf (Fig. 3c, 4e). The expansion of the putative pharyngeal arch one population (cluster S5), which was enriched for *cyp26c1* and *irx1b* transcripts ^60^, was also confirmed in Gata5/6 morphants by RNA ISH (Fig. 3c, 4f). For other cranial-mesoderm populations, we observed either small increases in marker gene expression (sub-cluster S2 and S6) or expression domains (sub-cluster S3), or no obvious changes (sub-cluster S7), largely consistent with the cell composition changes in our single-cell experiments (Fig. 3c and Supplementary Fig. 4b).

To examine a direct, cell-autonomous role for Gata5/6 in regulating cardiac and pharyngeal cell fate, transplantation experiments were performed. In these experiments, we assessed the ability of WT or Gata5/6-depleted cells isolated from donor embryos to contribute to cardiac and pharyngeal lineages. We used *Tg (nkx2.5: zsYellow)*^*fb7*^ embryos either uninjected or injected with Gata5/6 MO as donors, as this transgenic line marks both the heart and pharyngeal mesoderm derivatives ^56,61^. We observed that Gata5/6 deficient donor *Tg (nkx2.5: zsYellow)*^*fb7*^ cells contributed more than WT to the pharyngeal (9.35% VS 2.45%, *p*-value < 0.004), but not at all to the cardiac (0.36% VS 7.35%, *p*-value < 0.0002), mesoderm (Fig. 4g, h). Similar transplant experiments using *Tg (acta1: GFP)*^*zf13*^ as donors showed that Gata5/6 deficient cells, as compared to WT, favored a branchiomeric, but not trunk, muscle fate (Supplementary Fig. 4g, h). Together, this showed that loss of Gata5/6 biased cells to a pharyngeal mesoderm fate in a cell-autonomous manner, possibly at the expense of the cardiac lineage.

### Dosage effects of *gata5/6* on cardiac and pharyngeal lineage specification

At the time of this work being carried out, *gata6* mutants were unavailable, and it was uncertain if published *gata5* mutant alleles were true nulls (*s26*, *tm236a* ^14^. While *gata5* mutant and Gata5 morphant phenotypes are highly comparable ^14^, non-specific side effects have been ascribed to MOs ^62–64^. We therefore aimed to validate the scRNA-seq results in mutant embryos by generating novel null mutations in *gata5* and *gata6* using CRISPR-Cas9 mediated genome editing ^65^. Both a *gata5* mutant with a 29 bp deletion in the second exon and a *gata6* mutant with an 11bp deletion in the second exon were recovered (Supplementary Fig. 5.1a). Analysis of mutants revealed redundant functions for *gata5* and *gata6* in cardiac development, with double homozygous mutants lacking the expression of cardiac progenitor markers at 13 hpf and later heart formation (Supplementary Fig. 5.1 b and c), in full agreement with the cardiac defects reported in independently generated zebrafish *gata5/6* mutants ^6^. In addition, mutant embryos formed much smaller pectoral fins with morphological defects (Fig. 4a, Supplementary Fig. 5.1d). Overall, the cardiac phenotypes of *gata5*/*gata6* double homozygous mutants largely phenocopied those in Gata5/6 morphants (Supplementary Fig. 5.1 e and f), validating the MO usage for earlier stages of development when no obvious morphological phenotypes allowed for pre-selection of *gata5/6* mutants.

Analysis of *gata5/6* compound homozygous mutants confirmed the reduction of the cardiac and fin bud mesoderm and the expansion of the pharyngeal lineages observed in the morphant embryos (Fig. 4 and Supplementary Fig. 5.1). By analyzing all nine genotypes obtained from incrossing the *gata5+/-; gata6+/-* double heterozygous mutants, we further observed that the cardiac and pharyngeal phenotypes can be well explained by the reduction of the *gata5/6* gene dosage, with *gata5* playing a more prominent role (Fig. 5 and Supplementary Figs. 5.2 and 5.3). The robust dosage effect of *gata5/6* combined with the transplant experiments strongly indicated that the dosage of *gata5/6* is a principal determinant underlying cardiac and pharyngeal divergence.

**Fig. 5.**
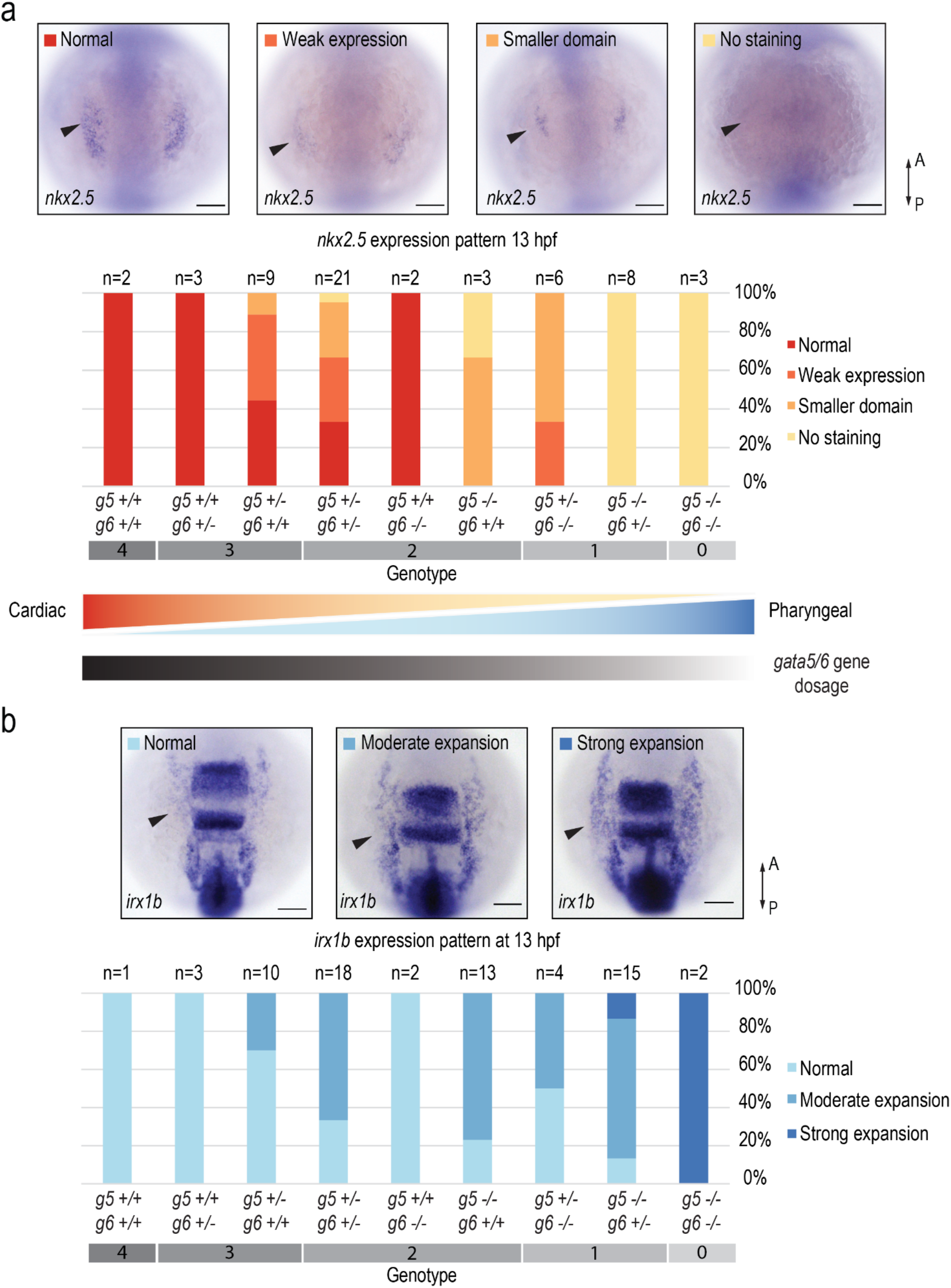
Cardiac and pharyngeal fate balance is controlled by *gata5/6* gene dosage. Gene expression analysis at 13 hpf of a marker gene for cardiac (S4: *nkx2.5*) (**a**) or pharyngeal (S5: *irx1b*) (**b**) lineage in nine different genotypes that were obtained from incrossing *gata5+/-*; *gata6+/-* double heterozygous mutants. Embryos are grouped and quantified into different categories according to their gene expression patterns. Representative embryos are shown in the top (**a**) or the bottom (**b**) panels. For each genotype, embryos that fall into different categories shown in bar plots with the number of embryos indicated above each bar.

### Gata4/5/6 promotes cardiac specification by antagonizing pharyngeal muscle identity in the tunicate *Ciona*

The invertebrate chordate *Ciona robusta* possesses a single *GATA4/5/6* homolog (*Gata.a*) ^8^, making it an ideal model to examine both the potential conservation and direct role for GATA4/5/6 function in cardiopharyngeal lineage divergence. Moreover, *Ciona* displays a simplified and stereotyped cardiopharyngeal lineage, whereby two multipotent progenitors on either side of the embryo each undergoes two asymmetric and oriented cell divisions to produce one first heart progenitor, one second heart progenitor and one pharyngeal muscle progenitor ^43,66^ (Fig. 6). Similar to zebrafish *gata5* and *gata6*, Ciona *Gata4/5/6* is expressed in migrating cardiopharyngeal progenitors ^8^, transiently down-regulated after their first division, and sequentially reactivated in the first and second heart precursor cells ^67^, a dynamic that is captured by trajectory reconstruction from scRNA-seq data ^68^. Thus, we used *Ciona* as a model to directly test whether Gata4/5/6 regulates the heart versus pharyngeal muscle fate decision in the cardiopharyngeal lineage.

**Fig. 6.**
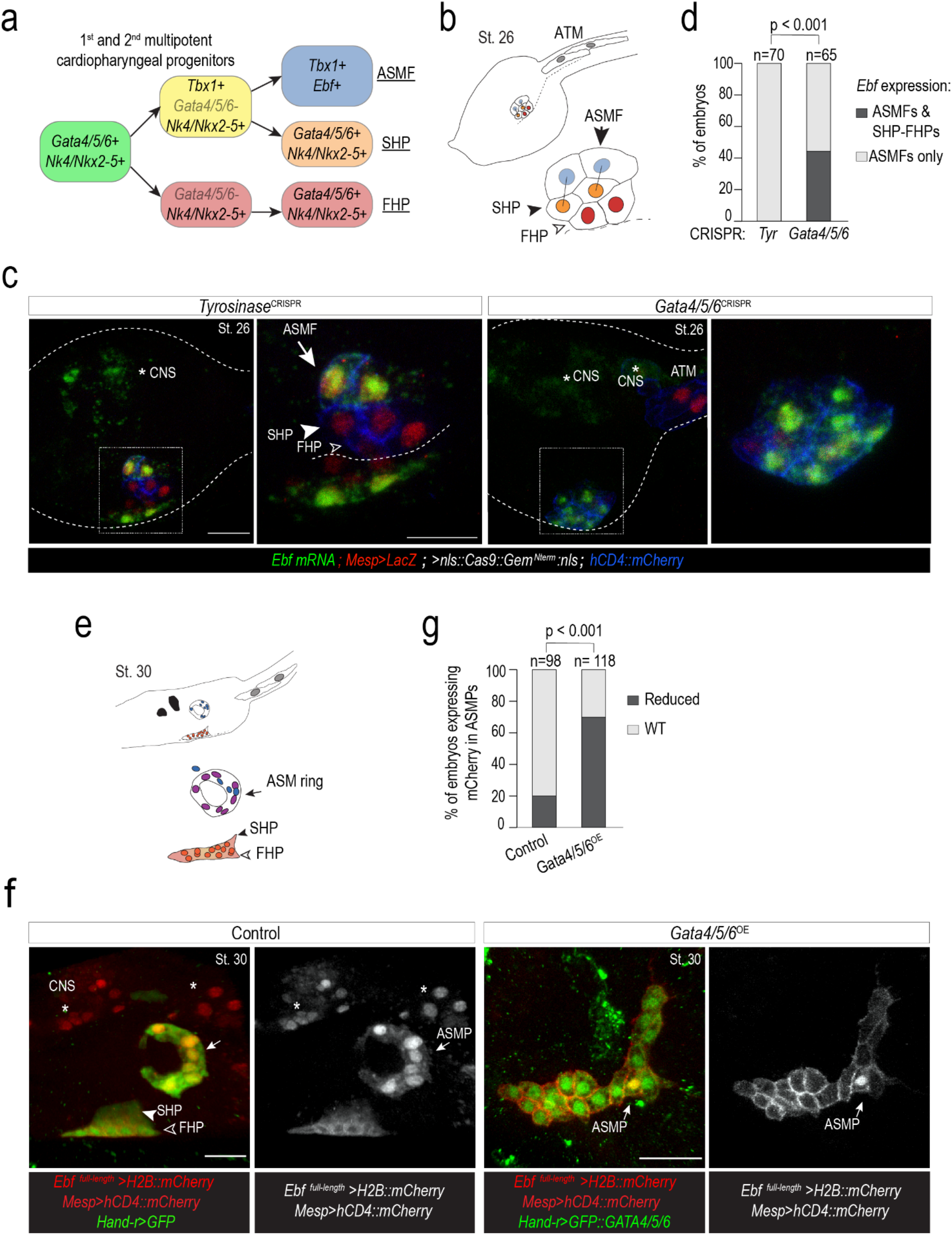
Gata4/5/6 is required to promote a cardiac and antagonize a pharyngeal muscle fate in Ciona. **a**, summarized cardiopharyngeal lineage in Ciona, showing expression status of indicated markers. Rounded squares represent cells and arrows represent cell divisions. Note the transient down-regulation of *Gata4/5/6* in early first heart progenitors (FHP, red) and *Tbx1+* 2^nd^ cardiopharyngeal progenitors (aka second trunk ventral cells, STVC, yellow), and subsequent reactivation in FHP and SHP (second heart progenitors (orange)), but not in ASMF (atrial siphon muscle founder cells (blue), or pharyngeal muscle progenitors), which activate *Ebf***. b**, Schematic representation of cardiopharyngeal lineage cells at Stage 26 ^102^. FHPs (red and open arrowheads), SHPs (orange and arrowheads), ASMF (light blue and arrows), anterior tail muscle (ATM); black bars link sister cells. **c**, Endogenous expression of *Ebf* visualized by ISH (green) in control (*Tyrosinase*^CRISPR^) and CRISPR/Cas9-mediated *GATA4/5/6* (*GATA4/5/6*^CRISPR^) knock-out embryos at stage 26. Nuclei of B7.5 lineage cells are labelled by *Mesp>nls::LacZ* and revealed with an anti-beta-galactosidase antibody (red). *Mesp*-driven hCD4::mCherry accumulates at the cell membrane as revealed by an anti-mCherry antibody (Blue). Scale bar = 15 μm or 10 μm (zoomed-in views for details in boxed regions). **d**, Proportions of embryos expressing *Ebf* transcripts in indicated cell-type progenitors. ASMF only (Tyr: 100%, n = 70), both ASMF and SHP-FHP (Tyr: 0%, *GATA4/5/6*^CRISPR^ : 43% ± 3% SE; n = 65); **e**, Schematic representation of cardiopharyngeal lineage cells at Stage 30 ^102^. FHPs: open arrowheads, SHPs: arrowheads, ASM ring: arrows. **f**, Target overexpression of *GATA4/5/6* in the cardiopharyngeal progenitor cells using a *Hand-r* enhancer. B7.5 lineage cells: red; GFP was revealed with an anti-GFP (green) and H2B::mCherry with an anti-mCherry antibody (red). Scale bar = 15 μm. ASMP: atrial siphon muscle precursor cells. **g**, Proportions of embryos expressing normal and reduced levels of *Ebf* in the ASMPs. Control: 20% (n = 98), *GATA4/5/6*^OE^: 68% ± 3% SE (n = 118) reduced *Ebf* expression in ASMPs Statistical analysis using a Fisher’s exact test showed all comparisons with controls to be significant (*p* < 0.01); “n” is the total number of individuals scored per condition. Note that only one side of the embryo is imaged. White asterisks refer to the central nervous system (CNS).

Using previously established reagents ^69^, we targeted CRISPR/Cas9-mediated mutagenesis to the cardiopharyngeal lineage and assayed expression of *Ebf*, a pharyngeal muscle determinant. We focused on stage 26 embryos, where the first and second heart lineages have already been segregated from pharyngeal muscles (Fig. 6a). Control embryos (*Tyrosinase*^*CRISPR*^) exhibited a WT pattern of *Ebf* expression, restricted to the lateral pharyngeal muscle precursor cells, also known as atrial siphon muscle founder cells (ASMF; Fig. 6a-d). By contrast, >40% of *Gata4/5/6*^*CRISPR*^ embryos exhibited conspicuous ectopic *Ebf* expression throughout the cardiopharyngeal lineage (Fig. 6a-d). Conversely, cardiopharyngeal lineage-specific over-expression of a GFP::Gata4/5/6 fusion protein sufficed to inhibit expression of an *Ebf* reporter construct and proper migration of pharyngeal muscle progenitors in ~70% of stage 30 swimming larvae, which normally exhibit a typical ring of pharyngeal muscles around the atrial siphon placode (Fig. 6e-g; ^43^). As *Ebf* acts as a master regulator of pharyngeal muscle identity and is sufficient to inhibit the cardiac fate ^43,69–71^, this result indicates that *Gata4/5/6* is necessary to both promote cardiac and prevent ectopic pharyngeal muscle fate in the common cardiopharyngeal progenitors. This further suggests that function of *GATA4/5/6* family members in the divergence of the cardiopharyngeal lineage is conserved between vertebrates and tunicates and is thus likely to be an ancestral feature of cardiac GATA factors.

## Discussion

Our study highlights that as opposed to being strictly a cardiac TF, zebrafish *gata5/6* are broadly expressed in multipotent mesendodermal progenitors and subsequently regulated in distinct manners in different developmental lineages. Multiple TFs have been shown to share similar dynamics including key regulators of pluripotency/neural development (*Sox2*) ^72^ and hematopoiesis (*Gata2*) ^73^. In addition to the *de novo* expressed lineage-specific TFs, we show in the case of *GATA4/5/6* that the selective silencing or maintenance of broadly expressed early TFs may serve as another key mechanism of cell fate determination. This complex regulation of gene expression is likely characteristic of TFs that can both promote and repress related cell fates, directly or indirectly, and may represent a widely used strategy in early development.

Single-cell gene expression atlases of whole zebrafish embryos continue to provide novel insights into early zebrafish embryogenesis ^35,38^. However, previous whole-embryo datasets have naturally been dominated by ectoderm-derived cells of the nervous system and epidermis, leaving mesoderm and endoderm lineages relatively under-represented. For example, at a comparable stage (14 hpf), one study profiled 677 mesoderm (17.2% in total) and 57 endoderm (1.45% in total) cells, roughly five times less than the mesendoderm cells (n=3,445) included in our 13 hpf data ^35^. By specifically focusing on the *gata5:GFP+* mesendoderm population, our study covers many of the lineages that are underrepresented in published whole embryo datasets. By further sub-clustering and RNA ISH characterization, we provide a high-resolution map of the molecular signatures of many cardiac and pharyngeal lineages. Future analysis of the function and expression of genes that define these populations will be of great interest. More generally, this lineage-focused scRNA-seq approach provides a powerful means to broadly phenotype without the need or cost of sequencing large numbers of cells.

Our study demonstrates that GATA-mediated cardiac-pharyngeal divergence is evolutionarily conserved between vertebrate and tunicates. Evidence suggests that the SHF shares a common progenitor pool with the cranial-pharyngeal mesoderm ^74,75^. Consistent with the previously described phenotype that SHF progenitors remained intact in *Gata4/6* mutant mice, we observed that a putative SHF population still formed in a *gata5/6* loss of function context in zebrafish, despite the heartless phenotypes in both models ^7^. In mouse embryos, the early expression patterns of *Gata4/6* follow a similar trend of broad expression in lateral plate mesoderm and later restricted expression in the developing heart ^18,28,76,77^. Future experiments that address how expression of *gata5/6* is maintained in cardiac and turned off in pharyngeal mesoderm will yield novel mechanistic insights into how these lineages segregate during embryogenesis. In this context, subtle alterations in the expression of key regulators such as *gata5/*6 may represent an important means to drive morphological changes via modulating cell type proportions.

GATA4 has been shown to have pioneer activity that can engage condensed chromatin and promote an open chromatin state *in vitro* ^26^. In Ciona, putative GATA binding sites are enriched in non-coding elements that open specifically in multipotent cardiopharyngeal progenitors in response to FGF-MAPK signaling ^70^. It is thus possible that GATA4/5/6 contributes to opening a substantial fraction of the cardiopharyngeal-lineage-specific cis-regulatory elements that control cardiac gene expression. Whether GATA4/5/6 pioneer activity is required in the GATA-mediated cardiac-pharyngeal divergence remains to be seen.

Besides the known role of GATA4/5/6 as transcriptional activators that promote cardiac specification ^31,78–80^, our observation that GATA4/5/6 are crucial for preventing the ectopic expression of the pharyngeal program suggests a repressive role. Indeed, it has been shown that mouse GATA4 can suppress the expression of non-cardiac programs ^81^ and a distal intestinal program in the proximal intestine through the interaction with its cofactor FOG ^82^. It is likely that GATA4/5/6 directly inhibit the pharyngeal program through a similar mechanism, perhaps by engaging distinct co-factors. Intriguingly, GATA family TF binding sites are frequently found in validated silencers, with a prevalence almost comparable to that of well-known chromatin repressors EZH2 and SUZ12. ^83^. Further studies, that look into temporal changes in co-binding factors and target genes of GATA4/5/6, and how GATA4/5/6 may integrate key signaling pathways, are needed to expand the gene regulatory network that governs development of the cardiopharyngeal lineage. While our work focused on the divergence of cardiac and pharyngeal fate, it is likely that the proper regulation of zebrafish *gata5/6* is also important for other lineages derived from *gata5*-expressing progenitors that require further characterization.

Our work uncovers highly dynamic, lineage-specific transcriptional regulation of *gata5/6* in multiple mesendoderm lineages during zebrafish gastrulation. We show that *GATA4/5/6* family members play key, conserved, and early roles in cardiac versus pharyngeal fate separation. This implies that precisely controlled *GATA4/5/6* expression during cardiac and pharyngeal specification is an ancestral mechanism that likely impacts an unexpectedly large number of additional mesendodermal lineages. Taken together, this adds support to a model where repression of key lineage regulators, such as *GATA4/5/6*, is as essential a step in development as their activation.

## Acknowledgements

We would like to thank Alejandro Salazar, Elyjah Schimmens, Melissa Westaway, Scott Knox and Nicholas Bendo for excellent animal care and zebrafish facility maintenance. Members of the Scott and Wilson labs provided invaluable suggestions and feedback during this project, in particular Huayun Hou (comments and discussions on computational analyses and the manuscript); Lynda Wu (pilot experiments) and Daniel Wagner at USCF for his help with the STITCH tool. Transgenic lines were kindly provided by Caroline and Geoffrey Burns (*Tg(nkx2.5: ZsYellow)*^*fb7*^), Kenneth Poss (*TgBAC (gata5:GFP)*^*pd25*^ and *TgBAC(tcf21:NSL-EGFP)*^*pd41*^), and James Dowling (*Tg (acta1: GFP)*^*zf13*^). We would like to thank the Centre for Applied Genomics for DNA sequencing and assistance with single-cell RNA-seq, in particular Matt Coole, Karen Ho, and Sergio Pereira, who were supported in part by a Genome Canada Genomics Technology Platform grant. This work was supported in part by NSERC grant RGPIN-2019-07014 to MDW, NIH awards R01 HD096770 and R01 HL108643 to LC, Ontario Trillium Scholarship and Ted Rogers Centre for Heart Research Education Fund to MS, and Hospital for Sick Children Restracomp Studentship and Connaught International Scholarship to XY. MDW is supported by the Canada Research Chairs Program and an Early Researcher Award from the Ontario Ministry of Research and Innovation. Key funding support was provided by NSERC (RGPIN-2017-06502 to ICS), the Heart and Stroke Foundation of Canada (G-16-00013798 to ICS) and the CIHR (FRN 86663 to ICS and FRN 156318 to MDW and ICS).

## Methods

### Zebrafish lines husbandry

All the zebrafish lines were maintained under the guidance and approval of the Canadian Council on Animal Care and the Hospital for Sick Children Laboratory Animal Services. Embryos were maintained at 28.5 °C in embryo medium and harvested at appropriate stages as previously described ^84^. To harvest samples at 13 hpf, embryos were first raised at 28.5 °C to 10 hpf and transferred to 22 °C overnight to 13 hpf. Staging was performed under the dissecting scope by embryonic morphology. The following transgenic lines were used in this study: *TgBAC (gata5:GFP)*^*pd25*^, *TgBAC (nkx2.5:ZsYellow)*^*fb7*^, *Tg(gata1a:dsRed)*^*sd2*^, *Tg(acta1:EGFP)*^*zf13*^, *Tg(myl7:EGFP)*^*twu34*^, *TgBAC(tcf21:NSL-EGFP)*^*pd41*^.

### Zebrafish CRISPR/Cas9 mutagenesis

CRISPR/Cas9 genome editing technology was used to generate *gata5* and *gata6* mutant lines as previously described ^65^. A 29 bp deletion allele *gata5*^*hsc115*^ (starting at nucleotide 3403 within exon 2) was generated, which results in a premature stop codon at amino acid 233. An additional allele with a 7 bp deletion displayed an equivalent phenotype. The 11bp deletion allele *gata6*^*hsc116*^ (starting at nucleotide 3522 within exon 2) was isolated with a premature stop codon formed at amino acid 332. Alternatively, another allele with a 10bp deletion and 5bp insertion displayed an equivalent phenotype. *gata5* heterozygotes were crossed to *gata6* heterozygotes to generate the compound heterozygous *gata5*^*hsc115*^; *gata6*^*hsc116*^ lines. Compound heterozygotes were then crossed to various transgenic backgrounds, including *Tg(myl7:EGFP)*^*twu34*^, *Tg(gata1a: dsRed)*^*sd2*^, *Tg(nkx2.5:zsYellow)*^*fb7*^, to perform downstream phenotypic characterization. Primers used for generating gRNAs are provided in Supplementary Table 2.

### Genotyping

Genomic DNA was extracted by incubating embryos or adult fin clips in 50mM NaOH at 95 °C for 20 min. 1M Tris PH8.0 was added at 1/10 volume to neutralize the reaction before PCR reactions. The *gata5* or *gata6* mutant allele was identified separately by amplifying a ~150 bp sequence containing the indel locus from the genomic DNA extracted from embryos or adult fin-clips. Mutant, heterozygous, and wild-type embryos were distinguished through the sizes of the PCR products. The same PCR cycling condition was used for all genotyping: denaturing at 94 ℃ for 5 min, thermocycling (35 cycles) at 94 ℃ for 30 s, 60 ℃ for 30 s and 72 ℃ for 30s, final elongation at 72 ℃ for 10 min. Primers used for genotyping *gata5* mutants: forward primer 5’-GACATGCTGGATGACCTGCCG −3’; reverse primer 5’-GCGTTTCTGTGGTTTGATAAGAG −3’. Primers used for genotyping *gata6* mutants: 5’-CGCGATTTATTGACTGAGTG −3’, and reverse primer 5’-CAGAGAAAGTGTCCCGTGC −3’.

### Morpholino injections

Morpholino (MO) oligonucleotides were purchased from Gene Tools. MOs targeting the splicing site of *gata5* (ssGata5: 5’-TGTTAAGATTTTTACCTATACTGGA −3’) and the translation start site of *gata6* (ATGGata6: 5’-AGCTGTTATCACCCAGGTCCATCCA −3’) were used as previously described ^5^. Injection of 9 ng of ssGata5 MO together with 1.5 ng of tbGata6 MO at one-cell stage displayed a heartless phenotype in line with cardiac defects in *gata5*^*hsc115*^, *gata6*^*hsc116*^ compound homozygous mutants.

### Colorimetric *in-situ* hybridization

Digoxigenin (DIG)-labeled and Fluorescein (Flu)-labeled in-situ probes were synthesized using *in vitro* transcription with Flu/DIG RNA Labeling kits respectively (Roche,Cat#11175025910, Cat#11685619910). RNA *in-situ* hybridization was conducted according to the published protocol ^85^. Embryos synchronized to the right developmental stages were fixed in 4% paraformaldehyde and then stored in 100% MeOH at −20 ℃ until used. After rehydration in PBSTw, embryos older than 48hpf were permeabilized using proteinase K before incubated with DIG-labeled antisense RNA probes at 65 ℃ overnight. RNA-probe hybrids were detected by an alkaline phosphatase-conjugated antibody (Anti-Digoxigenin-AP or Anti-Fluorescein-AP, Fab Fragments, 1:5000, Roche, Cat# 11093274910, Cat#11426338910. Embryos were washed in MABT before stained with NBT/BCIP (Roche, Cat#11681451001). Stained embryos were cleared in BBA solution (2:1 Benzyl benzoate: Benzyl alcohol) and imaged under a Zeiss Axio Zoom.V16 Stereoscope. Each *in-situ* hybridization experiment was performed at least twice to identify a certain gene expression pattern in both WT and Gata5/6KD embryos (n > 15). Embryos in-crossed from *gata5/6* compound heterozygous mutants were dehydrated in 80% glycerol for imaging and genotyped after imaging.

The following probes used have been described previously: *myl7* ^86^*, nkx2.5* ^87^*, tbx20, mef2cb* ^88^*, hand2, gata5, gata6, gfp* ^89^*, fgf8a* ^90^*, tbx5a* ^91^. Other *in-situ* probes were synthesized using the primers below:, *nr2f1a* (5’-ATGGTAGTTAGCGTCTGGCG-3’, 5’-AAAGCTTGCCAAAGCGACTG-3’), *irx1b* (5’-ACAGAGTTCCTCGGGAGT-3’, 5’-CCGATGACGTATATGCTGTTGC-3’), *cyp26c1*(5’-GATGCCGTCTTACCGACAGT-3’, 5’-ATCGCAGTAATCGCTGGCTTG-3’), *cthrc1a* (5’-CTGCATTTCGGTAATGATGG-3’, 5’-TGGTGATGCTCATTTTGGAAGC-3’), *kazald2* (5’-ATGCTGGTTTCCGTATC-3’, 5’-CACGGTGAGCTAATATTTCTGGTT-3’), *foxd1* (5’-CATCGGGAAACCCGAGAG-3’, 5’-ATTAGCAGCCGCTAGTGCCACTGA-3’), *six3b* (5’-TCCCCGTCGTTTTGTCTCTG-3’, 5’-TGTCCGACGGTGCATCATAC-3’), *tbx1* (5’-AAGGAGCGCAGTGGATGAAG-3’, 5’-GTACATGTTGGCTGTGGTTG-3’), *aldh1a2* (5’-TCGGCAGGGGAAAAAAACCC-3’, 5’-TTTTTTTTTTTTTTTCAGAGGTAA-3’).

### Transplantation and statistical test

Transplantation was performed as previously described ^92^. Transgenic lines incrossed from *Tg (nkx2.5: zsYellow)*^*fb7*^ or *Tg (acta1: GFP)*^*zf13*^ were used as donors to indicated cell fates for heart and pharyngeal arteries, trunk, and facial muscle, respectively. At 4 hpf, around 20-30 cells taken from the animal cap of these transgenic embryos either uninjected or injected with Gata5/6 MOs were placed into the margin of wild-type host embryos. Host embryos were imaged and scored at 48 or 72 hpf for the contribution of donor cells to cell fates or regions of interest using a Zeiss Axio ZoomV16 Stereoscope. Fisher exact tests were performed (**: *p-* value < 0.01, ***: *p-*value < 0.001).

### Fluorescent *in situ* hybridization with immunofluorescence

*TgBAC (gata5: EGFP)*^*pd25*^ embryos at the right developmental stages were fixed in 4% paraformaldehyde at 4 ℃ overnight. Prehybridization, hybridization and the saline-sodium citrate (SSC) washes were performed at 65°C ^85^. Detection of Digoxygenin (DIG) labeled probes were performed using an anti-DIG antibody, conjugated with horse-radish peroxidase (POD) (Roche Cat#11207733910) 1: 5000 followed by incubation with 1:50 to 1: 100 TSA plus Cyanine 3 solution (Perkin Elmer). To detect two genes at a time, embryos were incubated with a probe mix containing DIG and Flu-labelled probes. Fluorescein labeled probes were detected using an anti-Flu-POD antibody (Roche Cat#11426346910) and deposition of 1:50 to 1: 100 TSA plus Cyanine 3 solution. DIG-labelled probes were detected using an anti-Dig-AP antibody, developed with FastBlue (Sigma, F3378) and NAMP (Sigma, N5000), and monitored under a dissecting scope. GFP was detected using primary antibody Rabbit anti-GFP 1:500 (Torrey Pines Biolabs) and secondary antibody 488 Goat anti Rabbit (Thermo Scientific, Cat# A-11008) 1:1000. Embryos were stained for DAPI 1:2000 before mounting in low melt agarose for imaging under a Nikon A1R Si Point Scanning Confocal at 20X and 40X magnification.

### Imaging and cell counting

Bright-field images and were taken using a Zeiss AXIO Zoom V16. Confocal microscopy was performed under a Nikon A1R Si Point Scanning Confocal at 40X magnification to determine the *tcf21+* cell number in *TgBAC(tcf21: NLS-EGFP)*^*pd41*^ embryos and to image heart morphology in *Tg(myl7:EGFP)*^*twu34*^ embryos. The numbers of *tcf21+* cells were counted using the Cell Counter plugin in Fiji (https://imagej.nih.gov/ij/index.html).

### Embryo dissociation and cell isolation

WT and Gata5/6KD *TgBAC (gata5: GFP)*^*pd25*^ embryos were synchronized and dechorionated with pronase at 1mg/ml (Sigma, Cat# 11459643001) at desired stages. For embryos at gastrulation stages (6 hpf, 8 hpf, 10 hpf), the dissociation was performed as previously described ^89^. Embryos at the early segmentation stage (13 hpf) were treated similarly but incubated with 500 ul of 0.25 Trypsin (Gibco, Cat#15090046)/ EDTA instead of TrpLE (Gibco, Cat#12604013) for dissociation. Fluorescence-activated cell sorting (FACS) was performed on a Sony SH800S Cell Sorter, MoFlo XDP or MoFlo Astrios with a 100 μm nozzle by the SickKids-UHN Flow and Mass Cytometry Facility. Live, GFP+ cells were isolated using a similar gating strategy as described previously ^89^. 20,000 to 50,000 *gata5:*GFP+ cells were obtained in one FACS experiment.

### Single-cell mRNA-seq on 10X Genomic platform

Single-cell cDNA libraries were prepared through the 10X Chromium Single Cell Gene Expression platform in the TCAG sequencing facility at Sickkids. After FACS, GFP+ cells were counted twice with a Countless Automated Cell Counter (Thermofisher) before adjusting to a suspension of 300-500 cells/ul in 10% FBS/DMEM. 3000-6000 cells per sample were loaded into the 10X Chromium machine, with roughly 30% of cells captured. After Gel Bead-In Emulsions (GEM) generation and expression library construction as per the manufacturer’s instructions, final sequencing was performed on an Illumina HiSeq 2500 platform (Rapid Run Mode). 1000 to 2000 cells were collected for each condition with a sequencing depth of 119 ± 11.9 thousand reads per cell, which were estimated to reach 78.6 ± 2.88 % saturation of the whole libraries (Supplementary Table 1). The median of unique read (unique molecular identifier, UMI) counts per cell varied from 5,845 to 10,366 in different libraries (Supplementary Table 1).

### Single-cell mRNA-seq data alignment, quality control, and clustering analysis

Cell Ranger (version 2.0.0) was used for initial alignment, filtering non-cellular barcodes, and UMI counting to generate cell gene-count matrices with default parameters. Reads were aligned to the Ensembl Zv10 Release 89 reference genome and counted with the Ensembl transcriptome (Release 89) with a *gfp* sequence manually added. Sequencing saturation was estimated in the Cell Ranger pipeline by downsampling sequencing depth in mean reads per cell and assessing library complexities subsequently. The cell gene-count matrices generated in the Cell Ranger pipeline were filtered to exclude low-quality cells (low number of genes detected, high percentage of mitochondria gene reads) or potential cell doublets (high number of UMI counts). Different thresholds were used for samples from different stages, as cell size, RNA amounts, and RNA capture efficiencies varied between samples. Our filtering thresholds were 6 hpf (genes > 1000, UMI < 25000, mitochondria percentage < 4%), 8 hpf (genes > 1000, UMI < 20000, mitochondria percentage < 4%), 10 hpf (genes > 500, UMI < 12000, mitochondria percentage < 4%), WT and Gata5/6 KD 13 hpf (genes > 1000, UMI < 30000, mitochondria percentage < 4%) which were more stringent than that used in the previous studies of zebrafish embryos ^35,38^. The filtered gene-count matrices were used as input of Seurat (version 3.0.0) for dimensional reduction, cell clustering, and marker gene identification of each cluster. In the Seurat pipeline, SCTransform ^93^ was selected for data normalization, and the effects of mitochondria gene counts and cell cycles were regressed out before identifying variable genes according to package instruction.

Only genes that were identified as variable genes in SCTransform and were detected in large than 10% of cells in a cluster were tested as signature genes. MAST ^94^ was used and genes with log|FC| > 0.25 and p_val < 0.01 were identified as signature genes. Signature genes that were well studied and annotated (known marker genes) or best represented the transcriptome differences between clusters were selected for heatmap plotting. The 4 WT samples were analyzed separately (Supplementary Fig. 2.1-2.4). For comparing 13 hpf WT and Gata5/6 KD samples, these two samples were merged directly for clustering analysis without special integration.

To verify the SHF-like cluster, marker genes of the SHF-like sub-cluster were first converted to their mouse orthologs using gprofiler2. GO enrichment was performed on the mouse orthologs (Binomial test, Supplementary Table 3) ^49,50^. To compare the genes in the SHF-like cluster with a mouse SHF dataset ^51^, the zebrafish orthologs (n = 221) of the mouse anterior SHF genes (E7.75 and E8.25, n = 212) determined by the authors through scRNA-seq were first obtained using gprofiler2 and intersected with marker genes identified from each sub-cluster separately.

### Pseudotime analysis

After clustering analysis within each sample, the 4 WT samples were merged with cluster identity kept and then normalized as one dataset using the SCTransform ^93^ method. Clusters of each lineage were extracted from this merged dataset based on marker gene expression and fate-mapping knowledge (Supplementary Table 4) and analyzed using the dynverse package ^95,96^ for building up a pseudotime trajectory. *gata5* and *gata6* expression in each trajectory was plotted with ggplot2. To compare the *gata5* expression dynamics between different lineage trajectories, the pseudotime at each time point was divided into 10 quantile bins and the mean *gata5* expression of each bin was calculated and plotted in a heatmap (Fig. 2e).

### Visualization of the developmental progression of *gata5* GFP+ cells

STITCH ^35^ was implemented to generate a forced-layout visualization as previously described. In short, the filtered cell x gene counts matrix from all WT time points were merged and normalized as the input together with the developmental timepoint ‘ID’ for each cell.

Variable genes were first identified for each time point. STITCH next constructed a single-cell graph by computing k-nearest neighbors for each time point and identifying neighboring cells in pairs of adjacent time points to stitch the graph with a pre-defined order (6 hpf, 8 hpf, 10 hpf, 13 hpf). Up to 200 nearest neighbors were identified and up to 20 nearest neighbors were retained for each cell (nodes) with edges linking them in between. The single-cell graph was then visualized using ForceAtlas2 layout in Gephi (https://gephi.org/). Nodes are colored based on either timepoints or major branches. The latter required a lineage tag for each cell obtained from the clustering information in Seurat.

### Ciona experiments

Gravid adults were obtained from M-Rep (San Diego, USA), and gametes obtained surgically and used for *in vitro* fertilization and electroporation as described ^97,98^. Lineage-specific loss-of-function by CRISPR/Cas9-mediated mutagenesis was performed as described in ^69^ with a *Mesp>nls::Cas9::Gem*^*Nter*^*::nls*, which contains the N-terminus of human Geminin and exhibited increased efficacy (Christiaen lab, unpublished observation), and *U6* promoter-driven single guide RNAs targeting *Gata4/5/6* or the control target *Tyrosinase* (a pigment cell-specific gene inactive in the cardiopharyngeal lineage) described and validated previously ^99^. Overexpression was achieved by subcloning the coding sequence of GFP::Gata4/5/6 fusion protein ^100^ downstream of the trunk ventral cell/cardiopharyngeal progenitor-specific minimal *Hand-r* enhancer ^101^. Cardiopharyngeal lineage cells were visualized with *Mesp>hCD4::mCherry* and *Mesp>nls::LacZ* reporters, which labels the membranes and nuclei, respectively. The atrial siphon muscle/pharyngeal muscle progenitors were identified with an *Ebf>H2B::mCherry* reporter ^67,70^ or by fluorescent *in situ* hybridization as described ^70^.

### Data Accession

The raw sequencing data and single-cell count matrices can be accessed at ArrayExpress (E-MTAB-9193).

#### Contributions

Conception and study design: M.S., X.Y., C.R., L.C., M.D.W., and I.C.S.; Acquisition of data: M.S., X.Y., C.R., M.L., and A.A.; Analysis and interpretation of data: X.Y., M.S., C.R., L.C., M.D.W., and I.C.S.; supervision of work: M.D.W. and I.C.S.; M.S., X.Y., L.C., M.D.W., and I.C.S. wrote the manuscript and all authors assisted with drafting and revision of the manuscript.

#### Corresponding authors

Correspondence should be sent to Ian C. Scott and Michael D. Wilson.

