## Supplemental Figures for "GATA4/5/6 family transcription factors are conserved determinants of cardiac versus pharyngeal mesoderm fate"

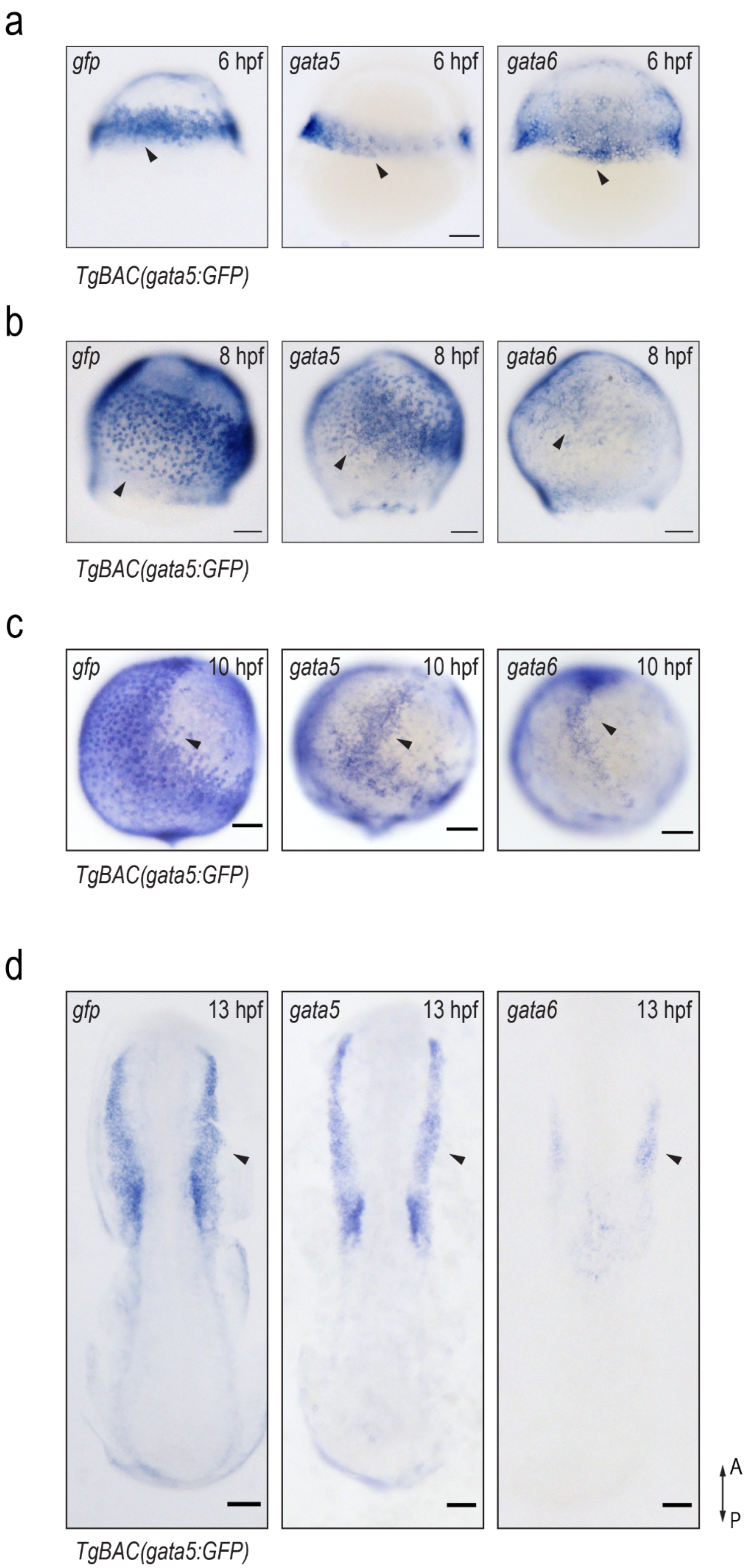
a
gfp
6 hpf
gata5
6 hpf
gata6
6 hpf
TgBAC(gata5:GFP)
b
gfp
8 hpf
gata5
8 hpf
gata6
8 hpf
TgBAC(gata5:GFP)
c
gfp
10 hpf
gata5
10 hpf
gata6
10 hpf
TgBAC(gata5:GFP)
d
gfp
13 hpf
gata5
13 hpf
gata6
13 hpf
A
P
TgBAC(gata5:GFP)

Supplementary Figure 1

Supplementary Figure 1.1

RNA ISH images of *gfp* in *TgBAC(gata5:GFP)* embryos, and *gata5* and *gata6* in WT embryos at 6 hpf (**a**), 8 hpf (**b**), 10 hpf (**c**) and 13 hpf (**d**). Arrowheads indicate the similar expression patterns of *gfp*, *gata5,* and *gata6*. A: anterior, P: posterior.

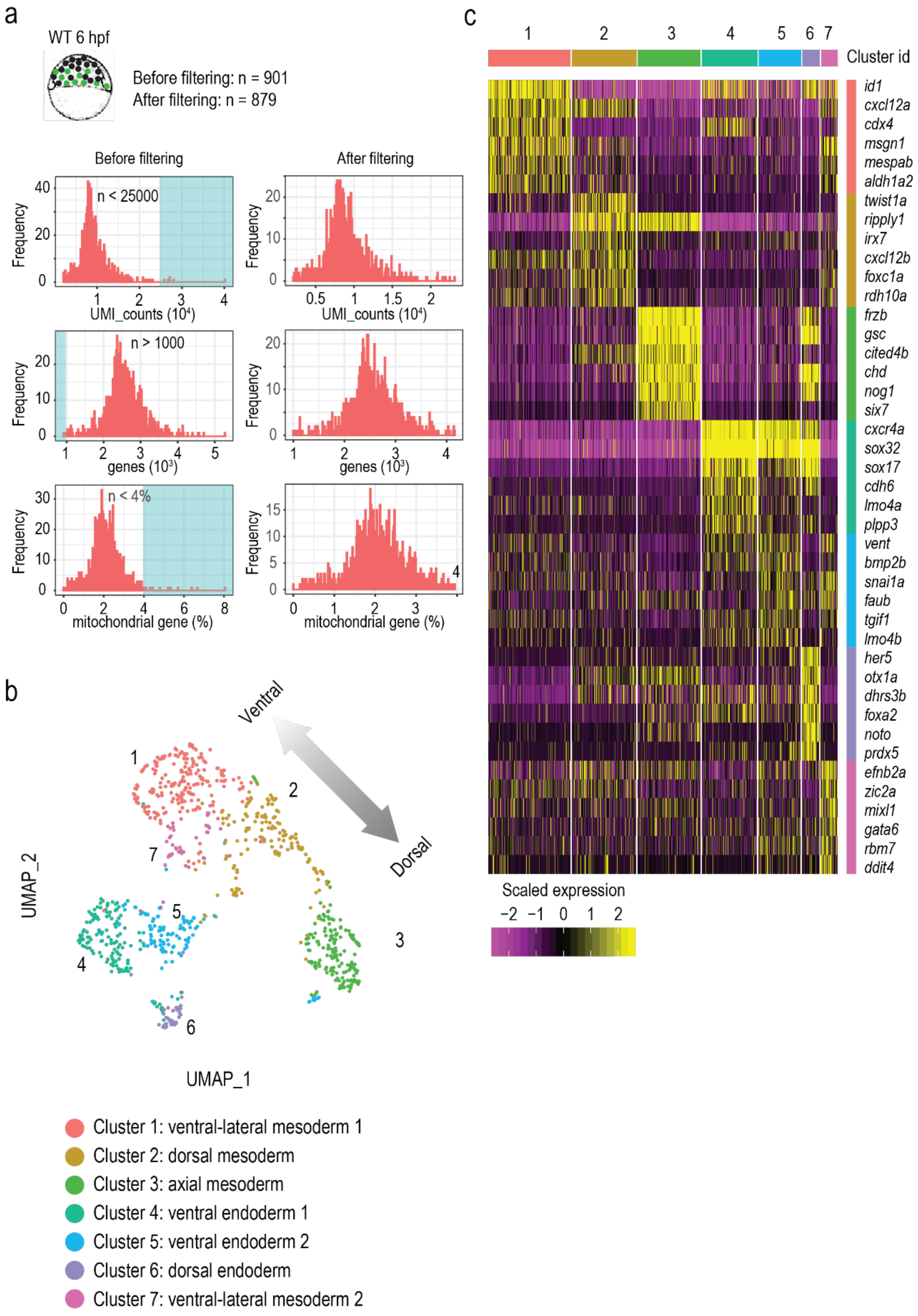
a
WT 6 hpf
Before filtering: n = 901
After filtering: n = 879
Before filtering
After filtering
Frequency
n < 25000
UMI_counts ($10^4$)
n > 1000
genes ($10^3$)
n < 4%
mitochondrial gene (%)
b
Ventral
Dorsal
UMAP_2
UMAP_1
Cluster 1: ventral-lateral mesoderm 1
Cluster 2: dorsal mesoderm
Cluster 3: axial mesoderm
Cluster 4: ventral endoderm 1
Cluster 5: ventral endoderm 2
Cluster 6: dorsal endoderm
Cluster 7: ventral-lateral mesoderm 2
c
Cluster id
id1
cxcl12a
cdx4
msgn1
mespab
aldh1a2
twist1a
ripply1
irx7
cxcl12b
foxc1a
rdh10a
frzb
gsc
cited4b
chd
nog1
six7
cxcr4a
sox32
sox17
cdh6
lmo4a
plpp3
vent
bmp2b
snai1a
faub
tgif1
lmo4b
her5
otx1a
dhrs3b
foxa2
noto
prdx5
efnb2a
zic2a
mixl1
gata6
rbm7
ddit4
Scaled expression
−2 −1 0 1 2

Supplementary Figure 1.2

(**a**) Histograms showing the distribution of UMI counts, numbers of genes detected, and percent of mitochondrial genes detected in the 6 hpf WT sample before and after filtering. Cyan shaded areas indicate cells that were filtered out to remove low-quality cells and potential doublets. (**b**) UMAP visualization of the 6 hpf WT sample coloured by cluster IDs (n = 879). (**c**) Heatmap showing the expression of top marker genes for each cluster in the 6 hpf WT samples.

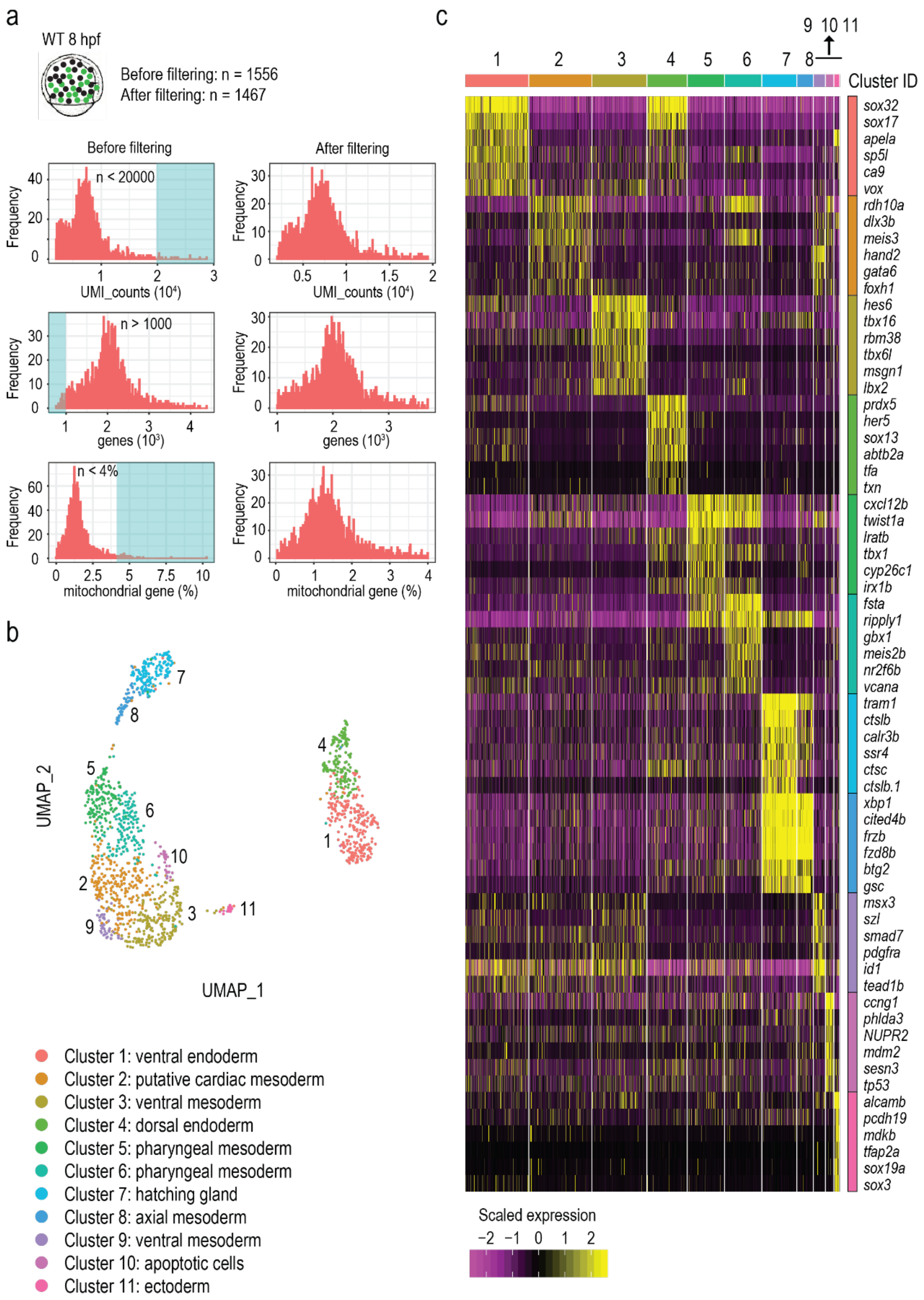
a
WT 8 hpf
Before filtering: n = 1556
After filtering: n = 1467
Before filtering
After filtering
n < 20000
Frequency
UMI_counts (10⁴)
n > 1000
genes (10³)
n < 4%
mitochondrial gene (%)
b
UMAP_1
UMAP_2
Cluster 1: ventral endoderm
Cluster 2: putative cardiac mesoderm
Cluster 3: ventral mesoderm
Cluster 4: dorsal endoderm
Cluster 5: pharyngeal mesoderm
Cluster 6: pharyngeal mesoderm
Cluster 7: hatching gland
Cluster 8: axial mesoderm
Cluster 9: ventral mesoderm
Cluster 10: apoptotic cells
Cluster 11: ectoderm
c
Cluster ID
sox32
sox17
apela
sp5l
ca9
vox
rdh10a
dlx3b
meis3
hand2
gata6
foxh1
hes6
tbx16
rbm38
tbx6l
msgn1
lbx2
prdx5
her5
sox13
abtb2a
tfa
txn
cxcl12b
twist1a
lratb
tbx1
cyp26c1
irx1b
fsta
ripply1
gbx1
meis2b
nr2f6b
vcana
tram1
ctslb
calr3b
ssr4
ctsc
ctslb.1
xbp1
cited4b
frzb
fzd8b
btg2
gsc
msx3
szl
smad7
pdgfra
id1
tead1b
ccng1
phlda3
NUPR2
mdm2
sesn3
tp53
alcamb
pcdh19
mdkb
tfap2a
sox19a
sox3
Scaled expression

Supplementary Figure 1.3

(**a**) Histograms showing the distribution of UMI counts, numbers of genes detected, and percent of mitochondria genes detected in the 8 hpf WT sample before and after filtering. Cyan shaded areas indicate cells that were filtered out to remove low-quality cells and potential doublets. (**b**) UMAP visualization of the 8 hpf WT sample coloured by cluster IDs (n = 1467). (**c**) Heatmap showing the expression of top marker genes for each cluster in the 8 hpf WT samples.

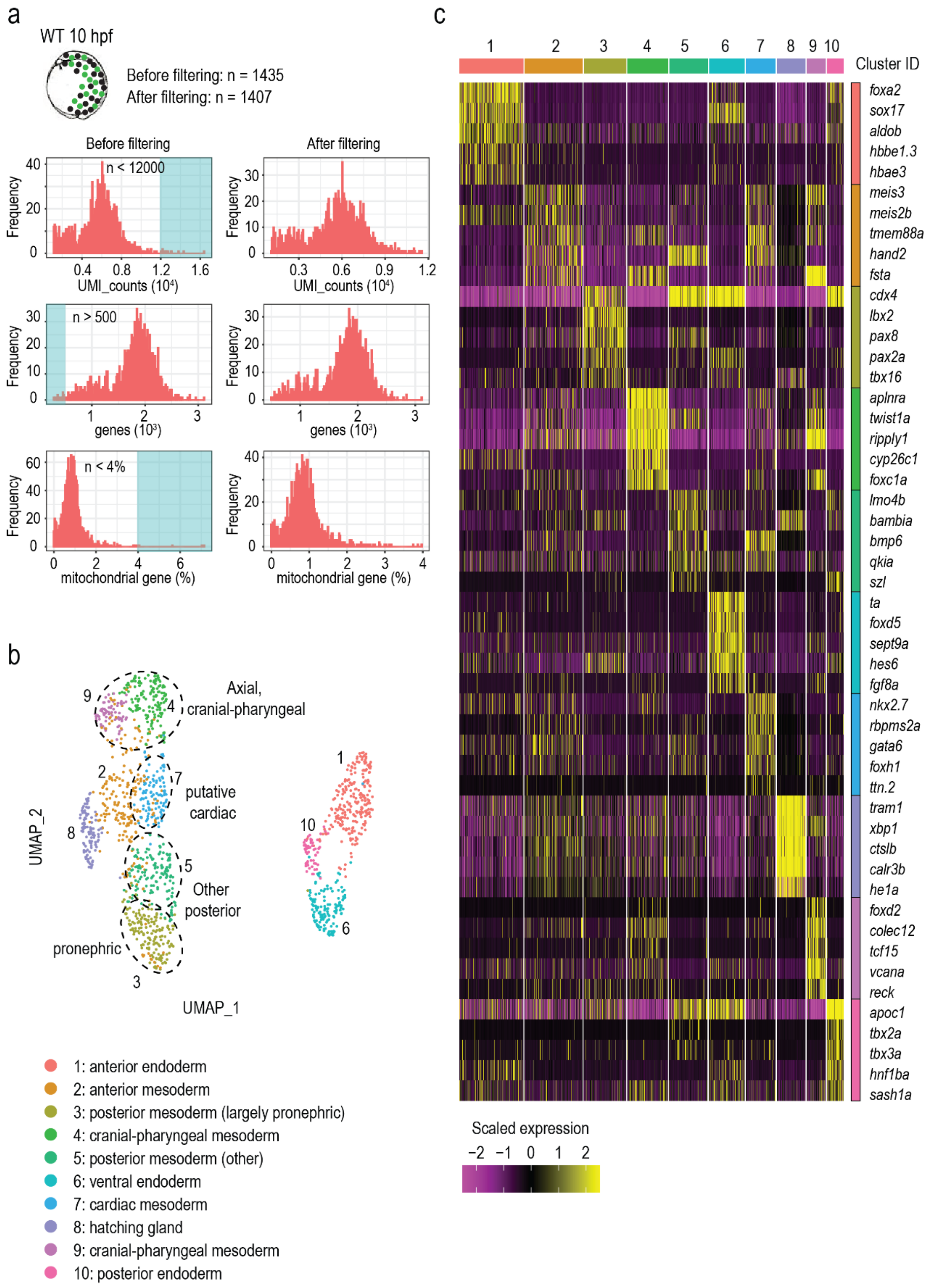

a
WT 10 hpf
Before filtering: n = 1435
After filtering: n = 1407
Before filtering
After filtering
n < 12000
Frequency
UMI_counts (10⁴)
n > 500
genes (10³)
n < 4%
mitochondrial gene (%)
b
Axial, cranial-pharyngeal
putative cardiac
Other posterior
pronephric
UMAP_1
UMAP_2
1: anterior endoderm
2: anterior mesoderm
3: posterior mesoderm (largely pronephric)
4: cranial-pharyngeal mesoderm
5: posterior mesoderm (other)
6: ventral endoderm
7: cardiac mesoderm
8: hatching gland
9: cranial-pharyngeal mesoderm
10: posterior endoderm
c
Cluster ID
foxa2
sox17
aldob
hbbe1.3
hbae3
meis3
meis2b
tmem88a
hand2
fsta
cdx4
lbx2
pax8
pax2a
tbx16
aplnra
twist1a
ripply1
cyp26c1
foxc1a
lmo4b
bambia
bmp6
qkia
szl
ta
foxd5
sept9a
hes6
fgf8a
nkx2.7
rbpms2a
gata6
foxh1
ttn.2
tram1
xbp1
ctslb
calr3b
he1a
foxd2
colec12
tcf15
vcana
reck
apoc1
tbx2a
tbx3a
hnf1ba
sash1a
Scaled expression

Supplementary Figure 1.4

(**a**) Histograms showing the distribution of UMI counts, numbers of genes detected, and percent of mitochondria genes detected in the 10 hpf WT sample before and after filtering. Cyan shaded areas indicate cells that were filtered out to remove low-quality cells and potential doublets. (**b**) UMAP visualization of the 10 hpf WT sample coloured by cluster IDs (n = 1407). (**c**) Heatmap showing the expression of top marker genes for each cluster in the 10 hpf WT samples.

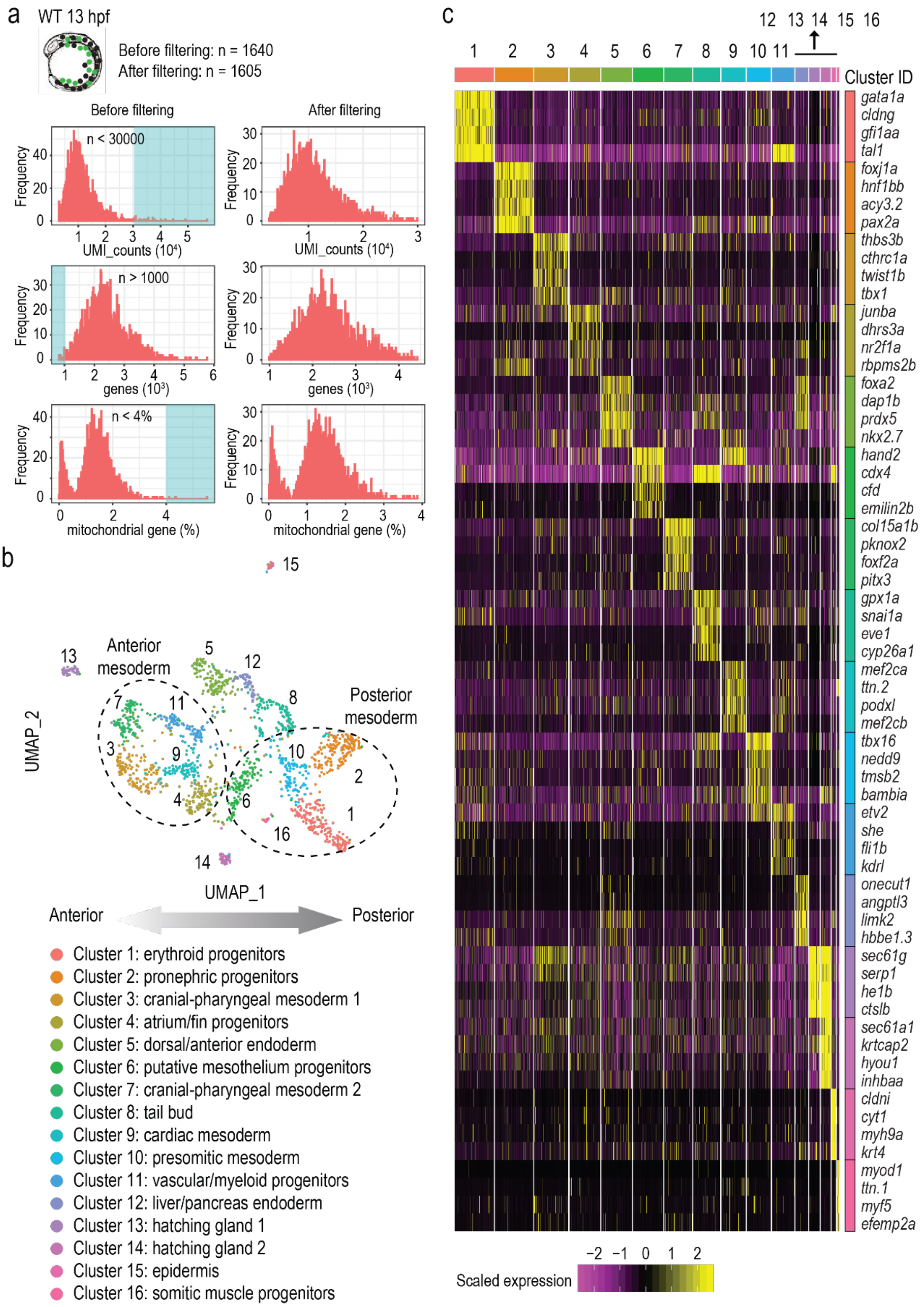
a
WT 13 hpf
Before filtering: n = 1640
After filtering: n = 1605
Before filtering
After filtering
n < 30000
UMI_counts (10⁴)
n > 1000
genes (10³)
n < 4%
mitochondrial gene (%)
Frequency
b
UMAP_2
UMAP_1
Anterior mesoderm
Posterior mesoderm
Anterior
Posterior
Cluster 1: erythroid progenitors
Cluster 2: pronephric progenitors
Cluster 3: cranial-pharyngeal mesoderm 1
Cluster 4: atrium/fin progenitors
Cluster 5: dorsal/anterior endoderm
Cluster 6: putative mesothelium progenitors
Cluster 7: cranial-pharyngeal mesoderm 2
Cluster 8: tail bud
Cluster 9: cardiac mesoderm
Cluster 10: presomitic mesoderm
Cluster 11: vascular/myeloid progenitors
Cluster 12: liver/pancreas endoderm
Cluster 13: hatching gland 1
Cluster 14: hatching gland 2
Cluster 15: epidermis
Cluster 16: somitic muscle progenitors
c
Cluster ID
gata1a
cldng
gfi1aa
tal1
foxj1a
hnf1bb
acy3.2
pax2a
thbs3b
cthrc1a
twist1b
tbx1
junba
dhrs3a
nr2f1a
rbpms2b
foxa2
dap1b
prdx5
nkx2.7
hand2
cdx4
cfd
emilin2b
col15a1b
pknox2
foxf2a
pitx3
gpx1a
snai1a
eve1
cyp26a1
mef2ca
ttn.2
podxl
mef2cb
tbx16
nedd9
tmsb2
bambia
etv2
she
fli1b
kdrl
onecut1
angptl3
limk2
hbbe1.3
sec61g
serp1
he1b
ctslb
sec61a1
krtcap2
hyou1
inhbaa
cldni
cyt1
myh9a
krt4
myod1
ttn.1
myf5
efemp2a
Scaled expression

Supplementary Figure 1.5

(**a**) Histograms showing the distribution of UMI counts, numbers of genes detected, and percent of mitochondria genes detected in the 13 hpf WT sample before and after filtering. Cyan shaded areas indicate cells that were filtered out to remove low-quality cells and potential doublets. (**b**) UMAP visualization of the 13 hpf WT sample coloured by cluster IDs (n = 1605). (**c**) Heatmap showing the expression of top marker genes for each cluster in the 13 hpf WT samples.

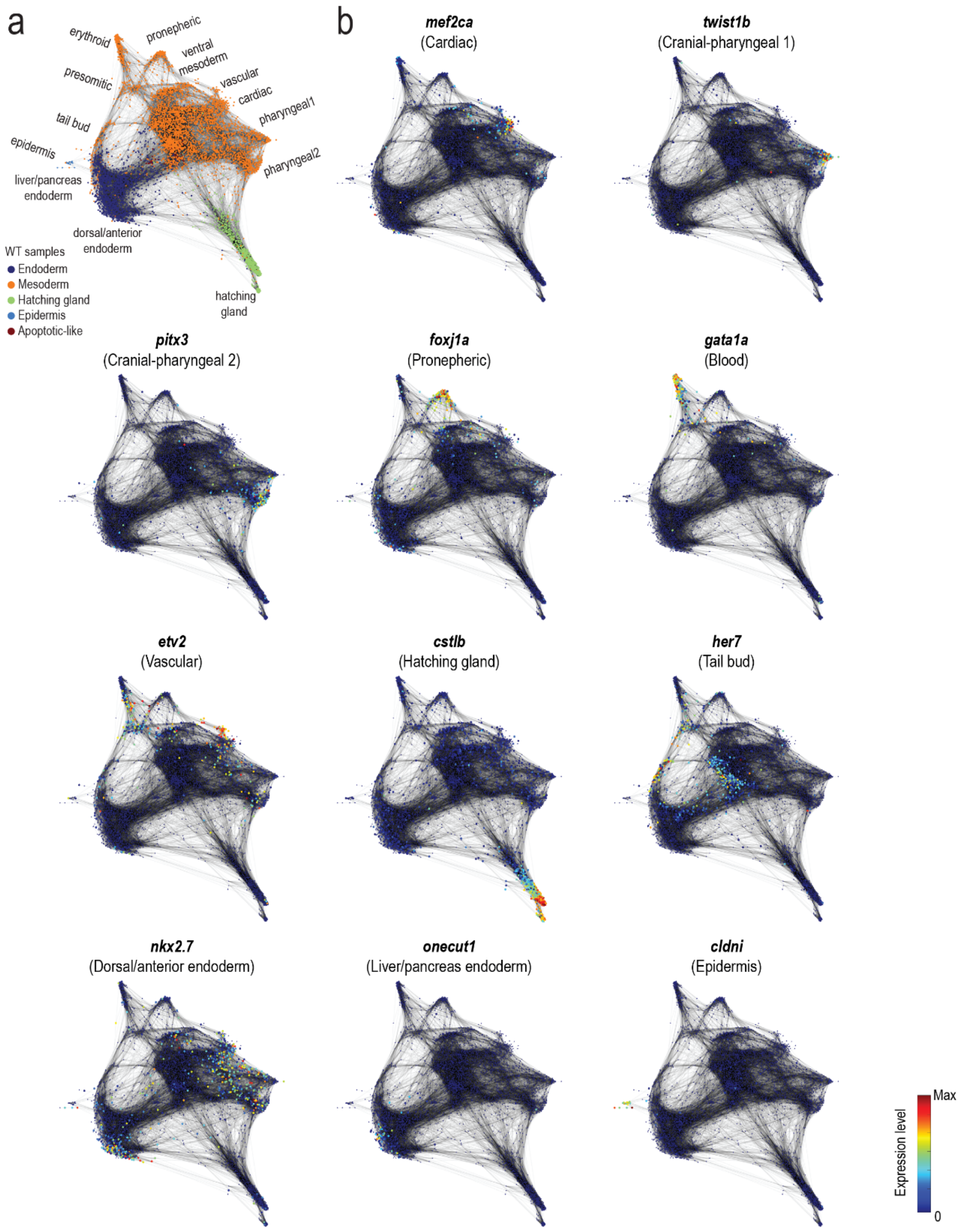
a
erythroid
pronepheric
ventral mesoderm
presomitic
vascular
cardiac
pharyngeal1
tail bud
epidermis
pharyngeal2
liver/pancreas endoderm
dorsal/anterior endoderm
WT samples
Endoderm
Mesoderm
Hatching gland
Epidermis
Apoptotic-like
hatching gland
b
mef2ca
(Cardiac)
twist1b
(Cranial-pharyngeal 1)
pitx3
(Cranial-pharyngeal 2)
foxj1a
(Pronepheric)
gata1a
(Blood)
etv2
(Vascular)
cstlb
(Hatching gland)
her7
(Tail bud)
nkx2.7
(Dorsal/anterior endoderm)
onecut1
(Liver/pancreas endoderm)
cldni
(Epidermis)
Expression level
Max
0

Supplementary Figure 1.6

(**a**) Force-directed graph showing the connection between all single cells from the four WT samples (6, 8, 10 and 13 hpf). Cells are colored based on their developmental origins (germ layers). (**b**) Marker gene expression of each lineage within the forced-directed graph.

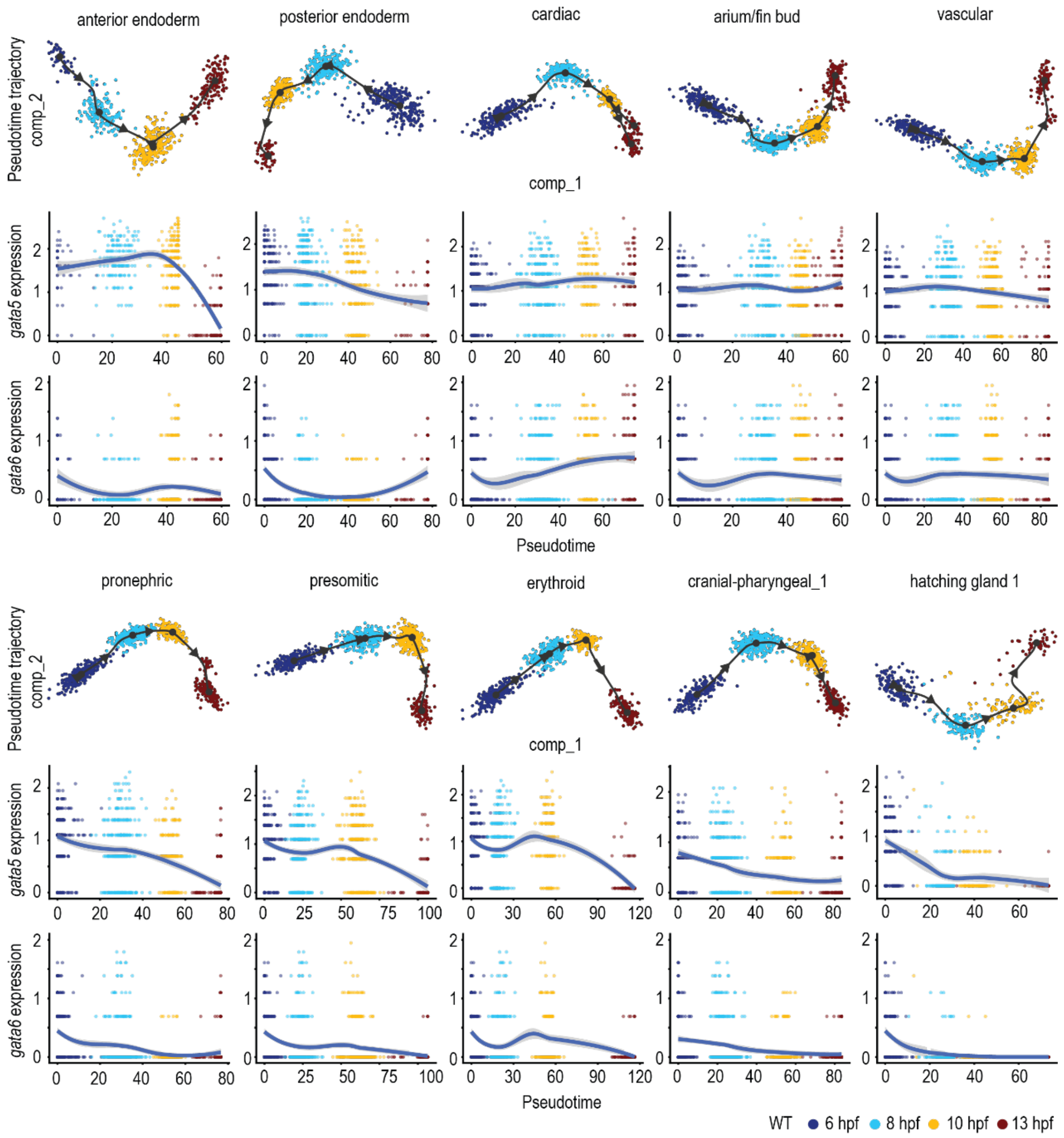

anterior endoderm
posterior endoderm
cardiac
arium/fin bud
vascular
Pseudotime trajectory comp_2
comp_1
gata5 expression
gata6 expression
Pseudotime
pronephric
presomitic
erythroid
cranial-pharyngeal_1
hatching gland 1
Pseudotime trajectory comp_2
comp_1
gata5 expression
gata6 expression
Pseudotime
WT 6 hpf 8 hpf 10 hpf 13 hpf

Supplementary Figure 1.7

Pseudotime developmental trajectories of the major lineages visualized in reduced dimensions (component_1 and component_2) and expression dynamics of *gata5* and *gata6* along each trajectory (x-axis, pseudotime; y-axis, normalized log expression level). In the gene expression scatter plots, lines show smoothed conditional means after local polynomial regression fitting (LOESS method) and shaded areas indicate standard errors.

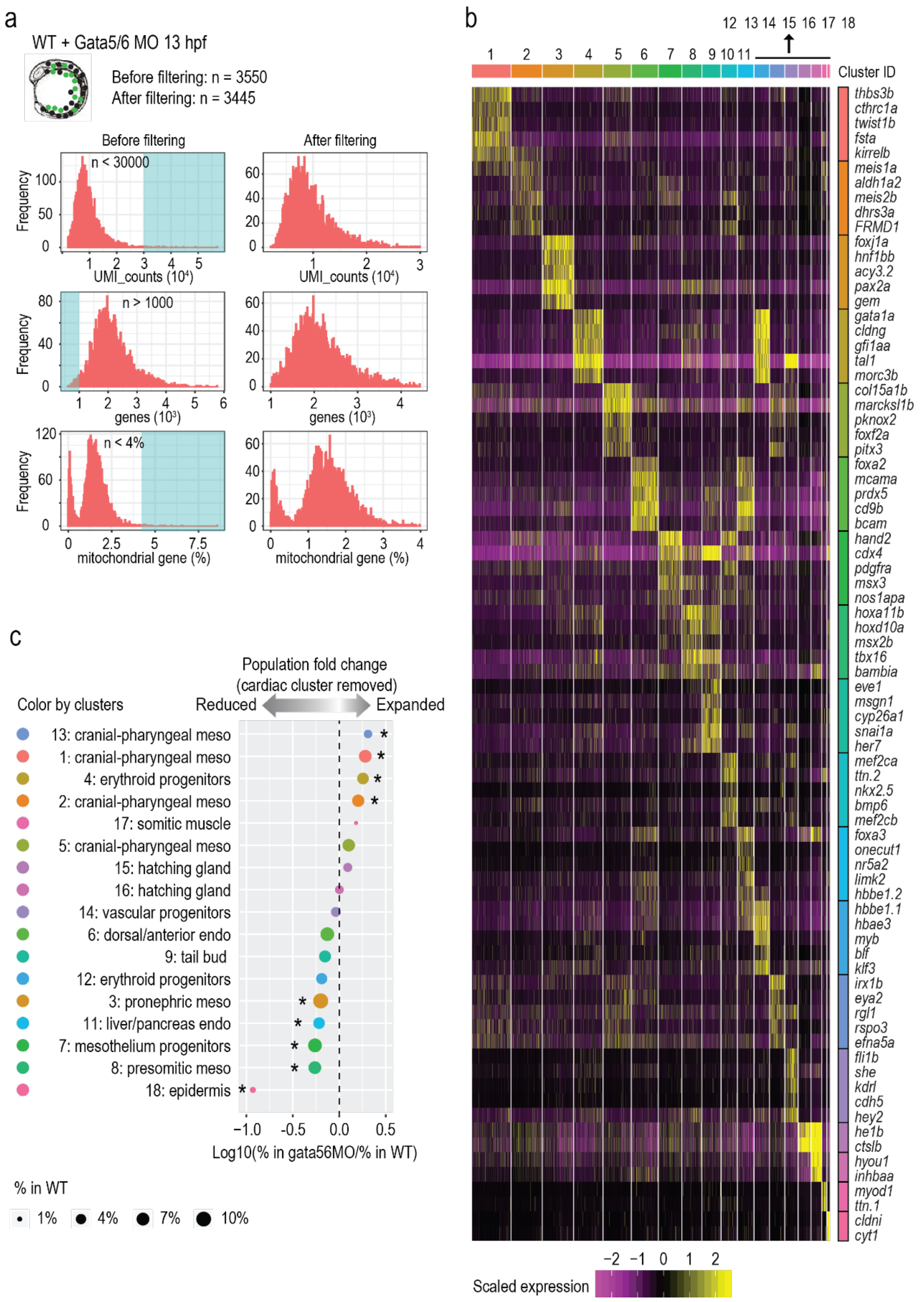
a
WT + Gata5/6 MO 13 hpf
Before filtering: n = 3550
After filtering: n = 3445
Before filtering
After filtering
Frequency
n < 30000
UMI_counts (10⁴)
n > 1000
genes (10³)
n < 4%
mitochondrial gene (%)
b
Cluster ID
thbs3b
cthrc1a
twist1b
fsta
kirrelb
meis1a
aldh1a2
meis2b
dhrs3a
FRMD1
foxj1a
hnf1bb
acy3.2
pax2a
gem
gata1a
cldng
gfi1aa
tal1
morc3b
col15a1b
marcksl1b
pknox2
foxf2a
pitx3
foxa2
mcama
prdx5
cd9b
bcam
hand2
cdx4
pdgfra
msx3
nos1apa
hoxa11b
hoxd10a
msx2b
tbx16
bambia
eve1
msgn1
cyp26a1
snai1a
her7
mef2ca
ttn.2
nkx2.5
bmp6
mef2cb
foxa3
onecut1
nr5a2
limk2
hbbe1.2
hbbe1.1
hbae3
myb
blf
klf3
irx1b
eya2
rgl1
rspo3
efna5a
fli1b
she
kdrl
cdh5
hey2
he1b
ctslb
hyou1
inhbaa
myod1
ttn.1
cldni
cyt1
Scaled expression
c
Population fold change
(cardiac cluster removed)
Color by clusters
Reduced
Expanded
13: cranial-pharyngeal meso
1: cranial-pharyngeal meso
4: erythroid progenitors
2: cranial-pharyngeal meso
17: somitic muscle
5: cranial-pharyngeal meso
15: hatching gland
16: hatching gland
14: vascular progenitors
6: dorsal/anterior endo
9: tail bud
12: erythroid progenitors
3: pronephric meso
11: liver/pancreas endo
7: mesothelium progenitors
8: presomitic meso
18: epidermis
Log10(% in gata56MO/% in WT)
% in WT
1%
4%
7%
10%

Supplementary Figure 2

(**a**) Histograms showing the distribution of UMI counts, numbers of genes detected, and percent of mitochondrial genes detected in WT and Gata5/6 MO combined samples at 13 hpf before and after filtering. Cyan shaded areas indicate cells that were filtered out to remove low-quality cells and potential doublets. (**b**). Heatmap showing the expression of top marker genes for each cluster in the 13 hpf WT and Gata5/6 MO combined samples (n = 3448). Five markers genes were plotted for most clusters except the last four clusters (hatching gland, somitic muscle, and epidermis) (**c**) Cell composition changes of each cluster between Gata5/6 KD and WT samples after the cardiac cluster was removed. Stars indicate significant differences (Fisher's exact test, Bonferroni correction, adjusted $p$-value < 0.05). Dot sizes show the percentage of each cluster within the whole WT population before cardiac cluster removal (total 13 hpf WT cells). Note that the cell composition changes kept the same trend for most clusters except that the reduction for liver endoderm became statistically significant due to less multiple test correction.

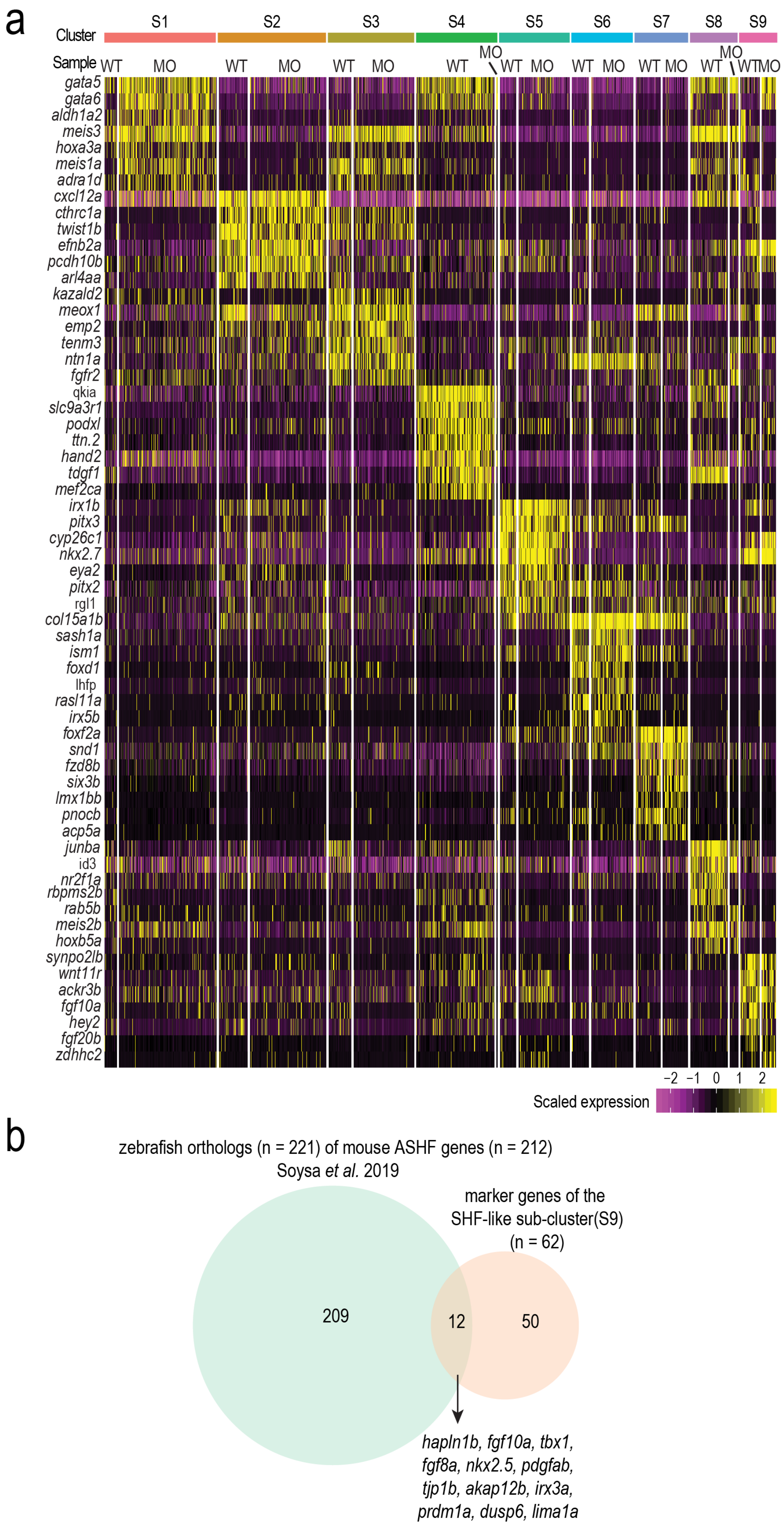
a
Cluster
S1
S2
S3
S4
S5
S6
S7
S8
S9
Sample
WT MO WT MO WT MO WT MO WT MO WT MO WT MO WT MO WT MO
gata5
gata6
aldh1a2
meis3
hoxa3a
meis1a
adra1d
cxcl12a
cthrc1a
twist1b
efnb2a
pcdh10b
arl4aa
kazald2
meox1
emp2
tenm3
ntn1a
fgfr2
qkia
slc9a3r1
podxl
ttn.2
hand2
tdgf1
mef2ca
irx1b
pitx3
cyp26c1
nkx2.7
eya2
pitx2
rgl1
col15a1b
sash1a
ism1
foxd1
lhfp
rasl11a
irx5b
foxf2a
snd1
fzd8b
six3b
lmx1bb
pnocb
acp5a
junba
id3
nr2f1a
rbpms2b
rab5b
meis2b
hoxb5a
synpo2lb
wnt11r
ackr3b
fgf10a
hey2
fgf20b
zdhhc2
−2 −1 0 1 2
Scaled expression
b
zebrafish orthologs (n = 221) of mouse ASHF genes (n = 212)
Soysa et al. 2019
marker genes of the
SHF-like sub-cluster(S9)
(n = 62)
209
12
50
hapln1b, fgf10a, tbx1,
fgf8a, nkx2.5, pdgfab,
tjp1b, akap12b, irx3a,
prdm1a, dusp6, lima1a

Supplementary Figure 3

(a) Heatmap showing the expression of top marker genes for each cardiac and pharyngeal cluster after sub-clustering, with WT and MO cells plotted separately. S1: posterior pharyngeal mesoderm, S2: cranial-pharyngeal mesoderm, S3: cranial-pharyngeal mesoderm, S4: cardiac progenitors, S5: first pharyngeal arch progenitors, S6: cranial-pharyngeal mesoderm, S7: cranial-pharyngeal mesoderm. (b) Venn diagram showing the overlap of markers genes in the SHF-like sub-cluster (S9, n = 62) and the zebrafish orthologs (n = 221) of the mouse anterior SHF genes (n = 212) identified through scRNA-seq (de Soysa et al., 2019).

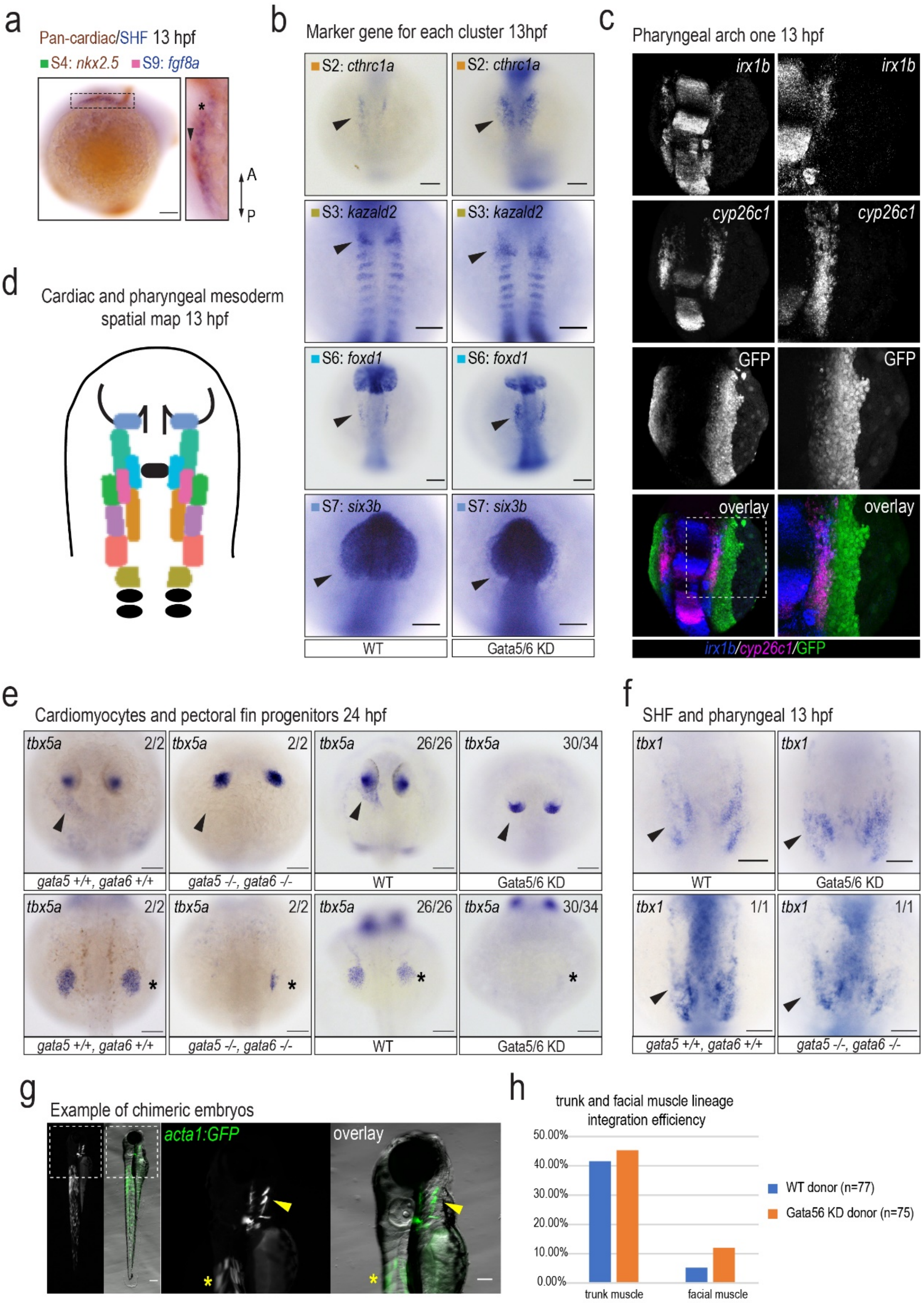
a
Pan-cardiac/SHF 13 hpf
S4: nkx2.5 S9: fgf8a
A
P
b
Marker gene for each cluster 13hpf
S2: cthrc1a
S2: cthrc1a
S3: kazald2
S3: kazald2
S6: foxd1
S6: foxd1
S7: six3b
S7: six3b
WT
Gata5/6 KD
c
Pharyngeal arch one 13 hpf
irx1b
irx1b
cyp26c1
cyp26c1
GFP
GFP
overlay
overlay
irx1b/cyp26c1/GFP
d
Cardiac and pharyngeal mesoderm spatial map 13 hpf
e
Cardiomyocytes and pectoral fin progenitors 24 hpf
tbx5a 2/2
tbx5a 2/2
tbx5a 26/26
tbx5a 30/34
gata5 +/+, gata6 +/+
gata5 -/-, gata6 -/-
WT
Gata5/6 KD
f
SHF and pharyngeal 13 hpf
tbx1
tbx1
WT
Gata5/6 KD
tbx1 1/1
tbx1 1/1
gata5 +/+, gata6 +/+
gata5 -/-, gata6 -/-
g
Example of chimeric embryos
acta1:GFP
overlay
h
trunk and facial muscle lineage integration efficiency
50.00%
40.00%
30.00%
20.00%
10.00%
0.00%
trunk muscle
facial muscle
WT donor (n=77)
Gata56 KD donor (n=75)

Supplementary Figure 4

(**a**) Double RNA ISH of *nkx2.5* (pan-cardiac) and *fgf8a* (SHF). (**b**) *In-situ* hybridization against marks for each sub-cluster in WT embryos and embryos injected with Gata5/6 morphants (Gata5/6 KD). Arrowheads indicate cranial-pharyngeal related expression domains of these genes that potentially overlap gata5:GFP+ marked regions A: anterior, P: posterior. (**c**) Double FISH against *irx1b* and *cyp26c1* (S5: pharyngeal arch one) together with GFP immunostaining on *TgBAC(gata5:EGFP)* embryos. (**d**) Schematic representation of the spatial organization of cardiac and pharyngeal mesoderm at 13 hpf. (**e**) RNA ISH against *tbx5a*at 24 hpf. Arrowheads indicate loss of *tbx5a* expression in the heart. Stars represent reduced *tbx5a* expression in pectoral fin bud progenitors. (**f**) RNA ISH against *tbx1* (S2, S5, S9). Arrowheads show that *tbx1* expression in anterior lateral mesoderm is largely unaffected upon loss of *gata5/6.* (**g**) Two representative chimeric embryos when *Tg (acta1:GFP)* transgenics were used as donors in transplant experiments, with arrowheads highlighting cells committed to a pharyngeal muscle fate and stars indicating a trunk muscle fate. (**h**) Quantification of the integration efficiency. SHF: second heart field.

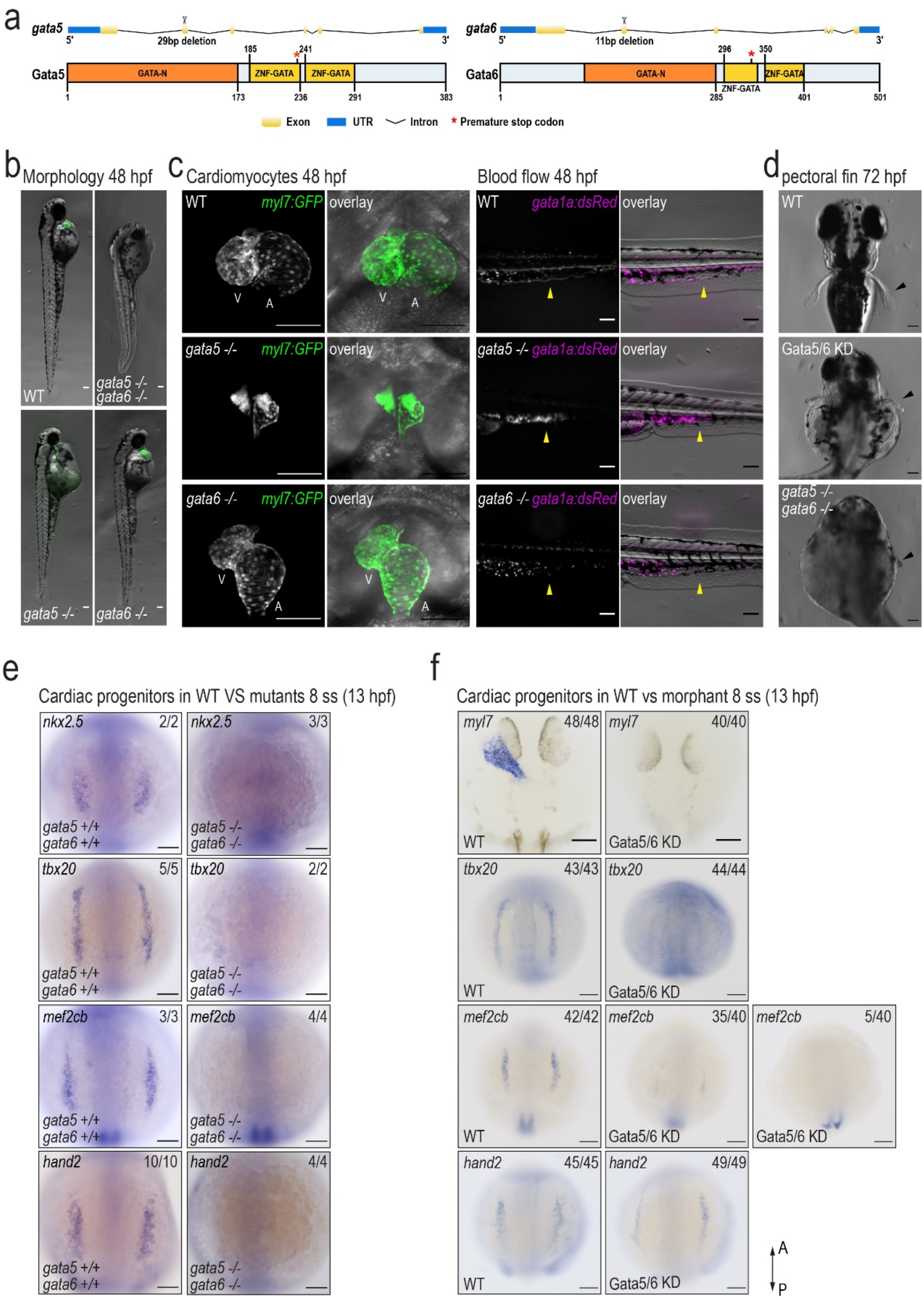

a
gata5
29bp deletion
Gata5
GATA-N
ZNF-GATA
ZNF-GATA
gata6
11bp deletion
Gata6
Exon
UTR
Intron
* Premature stop codon
b Morphology 48 hpf
WT
gata5 -/- gata6 -/-
gata5 -/-
gata6 -/-
c Cardiomyocytes 48 hpf
myl7:GFP
overlay
Blood flow 48 hpf
gata1a:dsRed
d pectoral fin 72 hpf
Gata5/6 KD
e
Cardiac progenitors in WT VS mutants 8 ss (13 hpf)
nkx2.5
tbx20
mef2cb
hand2
gata5 +/+ gata6 +/+
gata5 -/- gata6 -/-
f
Cardiac progenitors in WT vs morphant 8 ss (13 hpf)
myl7
A
P

Supplementary Figure 5.1

(**a**) Schematic representation of the *gata5* and *gata6* null-alleles generated through CRISPR/Cas9-mediated genome editing. A 29bp deletion was generated in exon2 of *gata5*. An 11bp deletion was introduced in exon2 of *gata6*. Stars indicate the premature stop-codon positions, at 233 amino acid in the first zinc finger domain of Gata5 protein and 332 amino acid in the first zinc finger domain of Gata6 protein. GATA-N: N-terminal GATA-type transcription activation domain; ZNF-GATA: Zince finger DNA binding domain. (**b**) Bright-field images of *Tg(myl7:EGFP)* at 48 hpf in a *gata5* null mutant, a *gata6* null mutant, a *gata5/gata6* double mutant and a sibling control from incrossing *gata5/gata6* compound heterozygous mutants. (**c**) Confocal images of *gata5*, *gata6* single mutants and their WT siblings in *Tg(myl7:EGFP)* and in *Tg(gata1a;dsRED)* backgrounds at 48 hpf. Yellow arrowheads indicate normal blood distribution in WT, blood pooling in *gata5-/-*, and impaired circulation in *gata6-/-* embryos. V: ventricle, A: atrium. (**d**) Bright field images of the pectoral fins in WT, Gata5/6 knockdown,and *gata5*, *gata6* compound homozygous mutant background at 72 hpf. Black arrowheads indicate the pectoral fins. (**e**) RNA ISH against known cardiac progenitor genes (*nkx2.5*, *tbx20*, *mef2cb*, and *hand2*) in *gata5/6* double mutants and WT siblings. (**f**) RNA ISH against known cardiac genes (*myl7, tbx20, hand2*) in WT embryos and embryos injected with Gata5/6 morpholinos (Gata5/6 KD). All scale bars represent 100 µm.

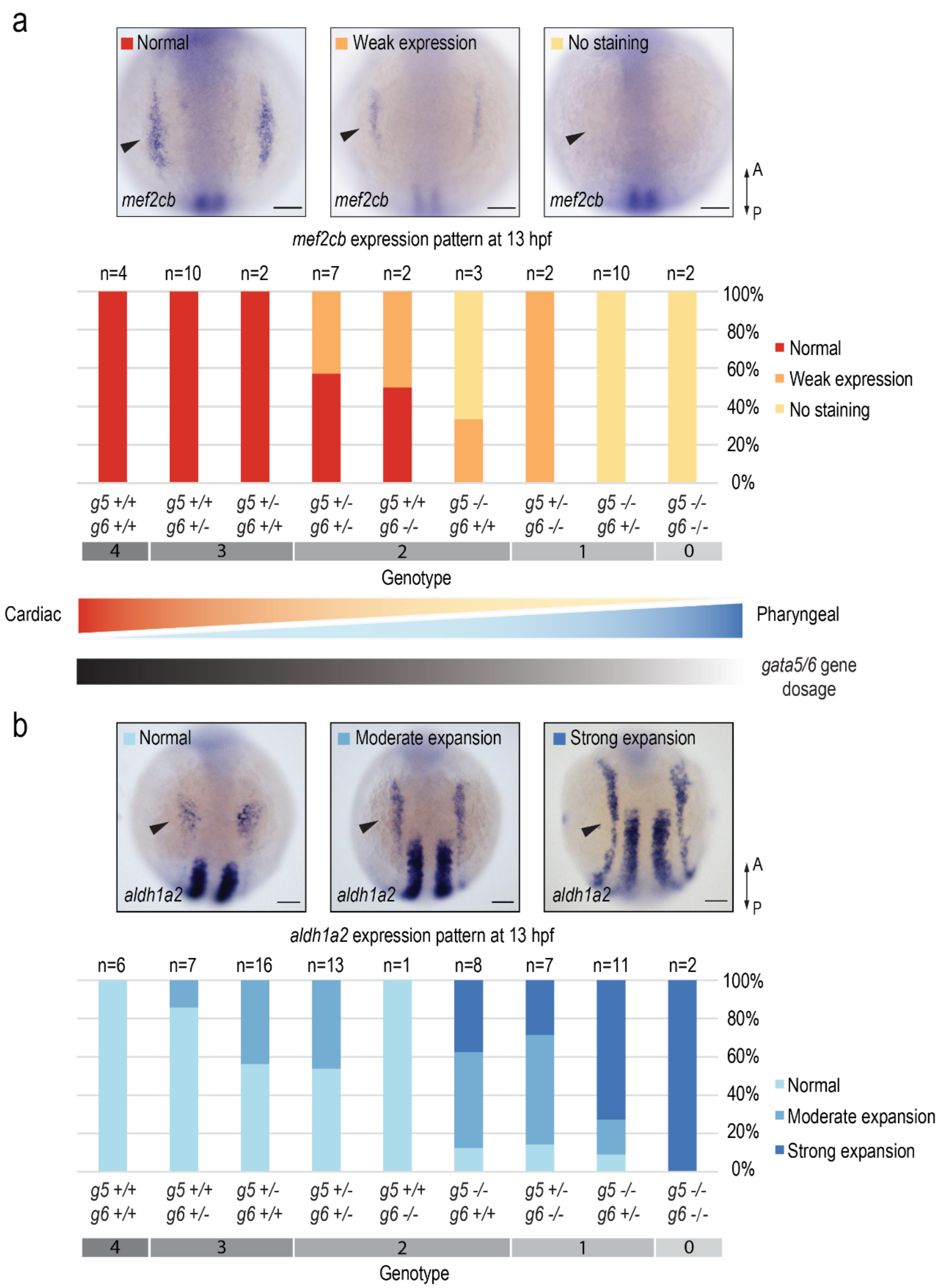
a
Normal
Weak expression
No staining
mef2cb
A
P
mef2cb expression pattern at 13 hpf
n=4
n=10
n=2
n=7
n=2
n=3
n=2
n=10
n=2
100%
80%
60%
40%
20%
0%
Normal
Weak expression
No staining
g5 +/+ g6 +/+
g5 +/+ g6 +/-
g5 +/- g6 +/+
g5 +/- g6 +/-
g5 +/+ g6 -/-
g5 -/- g6 +/+
g5 +/- g6 -/-
g5 -/- g6 +/-
g5 -/- g6 -/-
4
3
2
1
0
Genotype
Cardiac
Pharyngeal
gata5/6 gene dosage
b
Normal
Moderate expansion
Strong expansion
aldh1a2
A
P
aldh1a2 expression pattern at 13 hpf
n=6
n=7
n=16
n=13
n=1
n=8
n=7
n=11
n=2
100%
80%
60%
40%
20%
0%
Normal
Moderate expansion
Strong expansion
g5 +/+ g6 +/+
g5 +/+ g6 +/-
g5 +/- g6 +/+
g5 +/- g6 +/-
g5 +/+ g6 -/-
g5 -/- g6 +/+
g5 +/- g6 -/-
g5 -/- g6 +/-
g5 -/- g6 -/-
4
3
2
1
0
Genotype

Supplementary Figure 5.2

Gene expression analysis of a marker gene for the cardiac (S4: *mef2cb*) (**a**) or pharyngeal (S1: *aldh1a2*) (**b**) lineage in nine different genotypes that were obtained from incrossing *gata5+/- gata6+/-* double heterogenous mutants. Embryos are grouped into different categories according to their gene expression patterns. Representative embryos are shown in the top (**a**) or bottom (**b**) panels. For each genotype, embryos that fall into different categories are counted and shown in the bar plots. As the *gata5/6* dosage goes down, the cardiac progenitor population decreases with a concurrent expansion of the pharyngeal population.

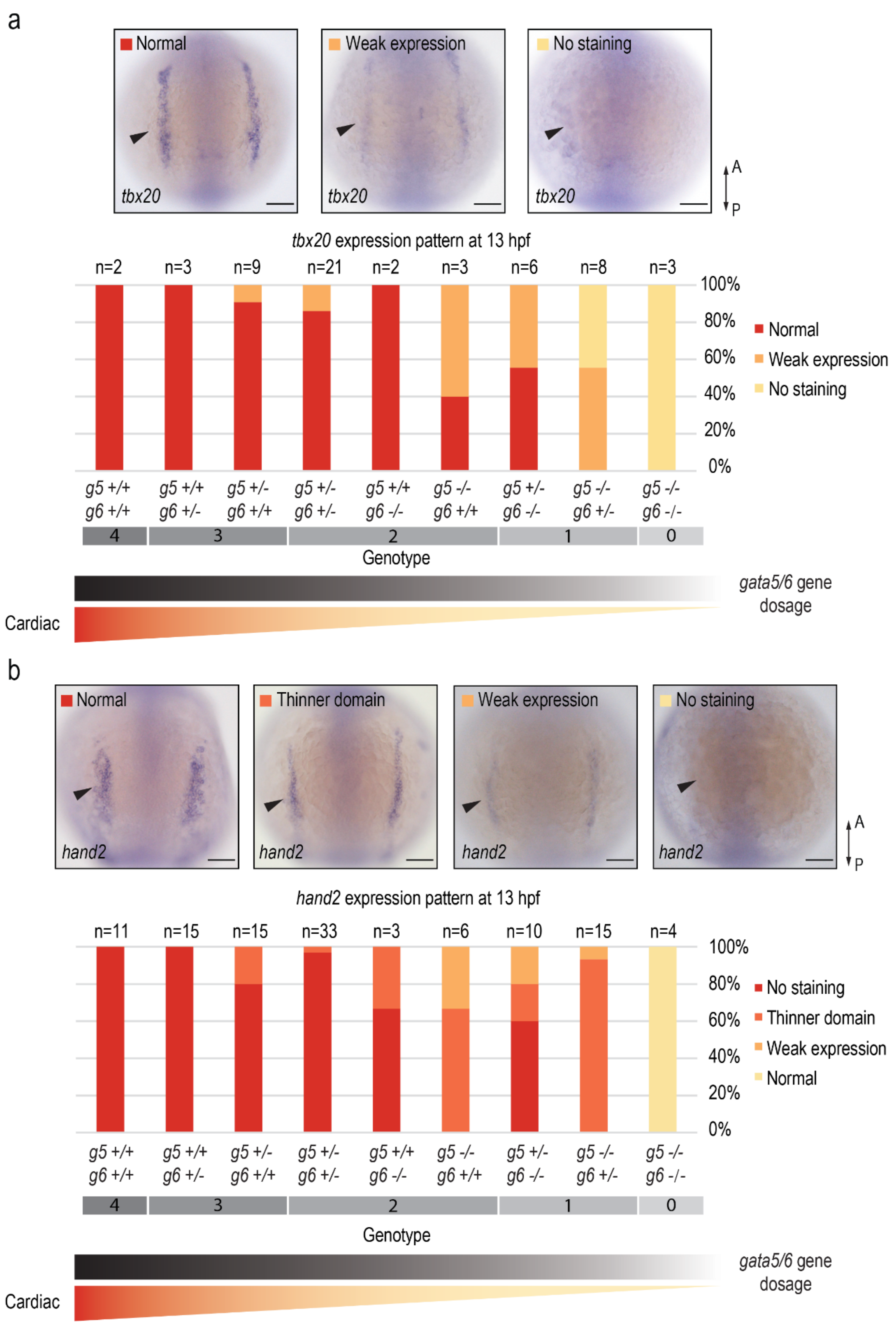
a
Normal
Weak expression
No staining
tbx20
A
P
tbx20 expression pattern at 13 hpf
n=2
n=3
n=9
n=21
n=2
n=3
n=6
n=8
n=3
100%
80%
60%
40%
20%
0%
Normal
Weak expression
No staining
g5 +/+ g6 +/+
g5 +/+ g6 +/-
g5 +/- g6 +/+
g5 +/- g6 +/-
g5 +/+ g6 -/-
g5 -/- g6 +/+
g5 +/- g6 -/-
g5 -/- g6 +/-
g5 -/- g6 -/-
4
3
2
1
0
Genotype
gata5/6 gene dosage
Cardiac
b
Normal
Thinner domain
Weak expression
No staining
hand2
A
P
hand2 expression pattern at 13 hpf
n=11
n=15
n=15
n=33
n=3
n=6
n=10
n=15
n=4
100%
80%
60%
40%
20%
0%
No staining
Thinner domain
Weak expression
Normal
g5 +/+ g6 +/+
g5 +/+ g6 +/-
g5 +/- g6 +/+
g5 +/- g6 +/-
g5 +/+ g6 -/-
g5 -/- g6 +/+
g5 +/- g6 -/-
g5 -/- g6 +/-
g5 -/- g6 -/-
4
3
2
1
0
Genotype
gata5/6 gene dosage
Cardiac

Supplementary Figure 5.3

Gene expression analysis of marker genes for the cardiac (S4: tbx20 (**a**) *and* hand2 (**b**)) in nine different genotypes that were obtained from incrossing *gata5+/- gata6+/-* double heterogenous mutants. Embryos are grouped into different categories according to their gene expression patterns. Representative embryos are shown in the top panels. For each genotype, embryos that fall into different categories are counted and shown in the bar plots. As the *gata5/6* dosage goes down, the cardiac progenitor population decreases.
